# AutoHiC: a deep-learning method for automatic and accurate chromosome-level genome assembly

**DOI:** 10.1101/2023.08.27.555031

**Authors:** Zijie Jiang, Zhixiang Peng, Yongjiang Luo, Lingzi Bie, Yi Wang

## Abstract

An accurate genome at the chromosome level is the key to unraveling the mysteries of gene function and unlocking the mechanisms of disease. Irrespective of the sequencing methodology adopted, Hi-C aided scaffolding serves as a principal avenue for generating genome assemblies at the chromosomal level. However, the results of such scaffolding are often flawed and require extensive manual refinement. In this paper, we introduce AutoHiC, an innovative deep learning-based tool designed to identify and rectify genome assembly errors. Diverging from conventional approaches, AutoHiC harnesses the power of high-dimensional Hi-C data to enhance genome continuity and accuracy through a fully automated workflow and iterative error correction mechanism. AutoHiC was trained on Hi-C data from more than 300 species (approximately five hundred thousand interaction maps) in DNA Zoo and NCBI. Its confusion matrix results show that the average error detection accuracy is over 90%, and the area under the precision-recall curve is close to 1, making it a powerful error detection capability. The benchmarking results demonstrate AutoHiC’s ability to substantially enhance genome continuity and significantly reduce error rates, providing a more reliable foundation for genomics research. Furthermore, AutoHiC generates comprehensive result reports, offering users insights into the assembly process and outcomes. In summary, AutoHiC represents a breakthrough in automated error detection and correction for genome assembly, effectively promoting more accurate and comprehensive genome assemblies.

## Introduction

The landscape of genomics research has undergone a remarkable transformation, unveiling the intricate tapestry of gene functionality and species evolution. Central to these breakthroughs is the pursuit of accurate^1^, chromosome-level genome sequences – an overarching goal that serves as the bedrock for unraveling the mysteries of biology and catalyzing the exploration of disease mechanisms.

Recent strides in genome assembly^2,3^ have been propelled by the emergence of long-read sequencing technologies, such as PacBio’s Single Molecule Real-Time^4^ (SMRT) sequencing and Oxford Nanopore Technologies^5^ (ONT). These technological marvels have shattered the constraints of traditional next-generation sequencing (NGS) methods^6^, offering read lengths that defy conventional limits. These factors have led to a spectacular surge in comprehensive and contiguous genome assemblies. However, even with these advancements, the elusive goal of achieving chromosome-scale assembly remains a challenge, underscoring the complexity of the task at hand.

In the intricate dance of genome assembly, Hi-C sequencing has arisen as a pivotal partner. This technique, an ingenious blend of proximity ligation and sequencing, promises to scaffold contigs into chromosome-scale assemblies by capitalizing on the higher density of Hi-C linkage pairs between adjacent contigs^7–12^. While a slew of tools, such as Lachesis^13^, 3D-DNA^14^, SALSA^15,16^, YaHS^17^, instaGRAAL^18^, EndHiC^19^, and Pin_hic^20^ have emerged to translate Hi-C data into chromosome-scale scaffolds, each harbors limitations and is susceptible to various influences. Some software requires the number of chromosomes to be specified in advance, but this is very difficult for the user. Moreover, the presence of errors in the output generated by these tools necessitates manual correction, extending the process and inviting human error. This dependence on manual intervention has cast a shadow over the dream of fully automated genome assembly, particularly when striving for chromosome-level precision with Hi-C data.

As a symphony of large-scale genomic research initiatives such as the Bird 10,000 Genomes (B10K) Project^21^, the Earth Bio Genome Project^22^ (EBP), and the i5k initiative^23^ take the global stage, the need to automate the process of high-quality chromosome-level genome assembly at scale has emerged as an urgent imperative. Nevertheless, with the ongoing expansion and increasing complexity of datasets, conventional assembly approaches come with challenges in terms of algorithmic precision and available human resources. This is where innovative assembly software, empowered by cutting-edge technologies such as artificial intelligence and deep learning, shines. This confluence promises to create a transformative force capable of untangling the intricate puzzle of genome splicing and assembly.

Furthermore, deep learning has come to play an increasingly pivotal role in the life sciences^24–26^, significantly contributing to data analysis and processing. Transformers^27^, which are a type of attention-based architecture designed for long sequences, have made remarkable strides in language processing and have demonstrated applicability in other domains, including image analysis, gene expression, and protein folding. Despite the emergence of software such as DeepC^28^, EagleC^29^, VEHiCLE^30^, DeepLoop^31^, SnapHiC-D^32^, and hicGAN^33^, the full potential of Hi-C data in the detection of assembly errors remains underutilized. Large datasets provide a potential basis for deep learning to fully exploit Hi-C information.

Here, we present AutoHiC, a scalable and computationally efficient deep learning-based error correction method. Leveraging genome-wide chromatin interaction data from over five hundred thousand Hi-C images derived from Hi-C data from approximately 300 species, AutoHiC automates realize Hi-C assembly error correction, significantly improving genome assembly continuity and accuracy. We demonstrate the feasibility of the AutoHiC recognition and error correction algorithm by comparing the interaction heatmaps before and after adjustment. From the continuity comparison results, it was found that AutoHiC can significantly improve genome continuity compared to other software. Moreover, to more accurately reflect the actual situation of the genome, we performed genome accuracy tests on the T2T genomes, and the results showed that AutoHiC can improve the accuracy of genomes. Finally, to prove the universality of AutoHiC, we tested it in several species, including those with large genomes, a large number of chromosomes, and polyploidy. The results showed that AutoHiC can be well applied to different situations.

In summary, AutoHiC harnesses the power of deep learning and Hi-C to automate chromosome-level genome assembly and advance scaffold assembly. By automatically identifying and correcting Hi-C assembly errors and achieving exceptional chromosome-level assembly results, AutoHiC supersedes laborious and error-prone manual adjustments, revolutionizing the final step of genome sequencing data to chromosome-level genome assembly with automation and efficiency. This groundbreaking advancement holds immense promise for advancing genomics research and deepening our understanding of genome three-dimensional structure and functionality.

## Results

### Overview of the AutoHiC pipeline

To provide a fully automated, deep learning-based approach to identifying and correcting misassembled genomes, we aimed to develop a genome tool using a deep learning-based approach. AutoHiC draws primarily on the lessons learned from manual error correction with assembly tools^34,35^ to provide a fully automated, deep learning-based approach to identifying and correcting misassembled genomes. This tool empowers users without computational backgrounds who are unfamiliar with Hi-C images to enhance genome quality and reduce the costs associated with manual adjustment of genome assembly. The overall workflow of AutoHiC is illustrated in Fig. 1a. A detailed description is available in the Methods section.

**Fig. 1:**
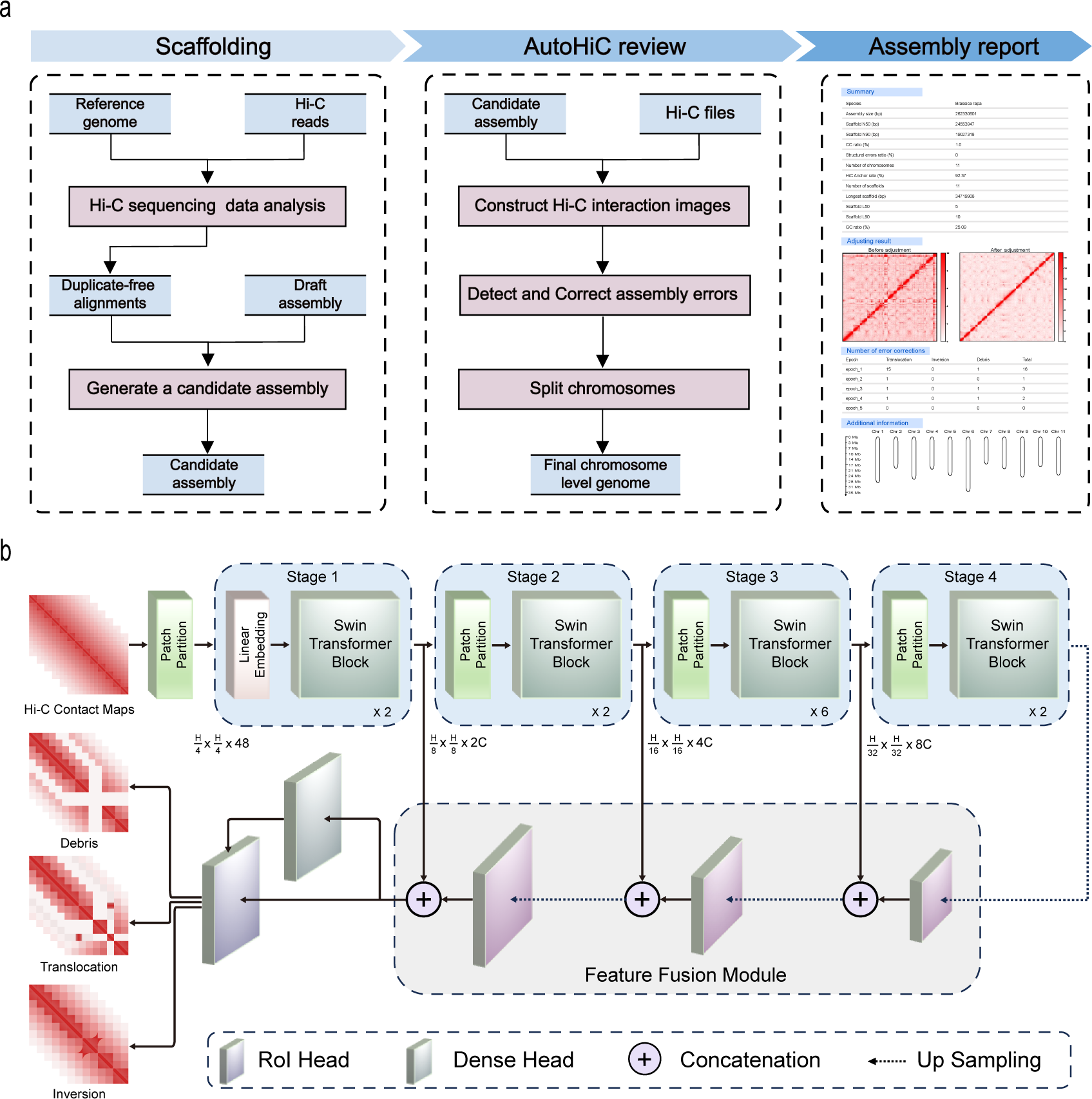
Overview of the AutoHiC framework. **a** AutoHiC runs processes and data flows. The AutoHiC assembly process is divided into three main steps. Users only need to prepare contigs and Hi-C reads the data. First, scaffold software is used to generate candidate genomes and global interaction files. Next, the AutoHiC model detects and iteratively corrects assembly errors and partitions chromosomes based on the interaction matrix. Finally, the assembly result report is generated. Data and steps are marked with different colors. **b** AutoHiC model description. The architecture of the Swin Transformer network employed in AutoHiC is depicted. The input is a three-channel interaction image, and the output is the type of error detected by the model and the corresponding location information. Different layers adopt different configurations, which are distinguished by colors, and the specific information subscripts are visible.

At a macroscopic level, AutoHiC operates in three stages. In the first stage, AutoHiC leverages Juicer^36^ and 3D-DNA to generate preliminary assembly results based on the existing contig and Hi-C reads. The outcomes of this step include scaffolds (uncorrected) and Hi-C interaction maps at various resolutions. In the second which is the most crucial step of the pipeline, AutoHiC incorporates an error correction module (Methods) to refine assembly quality, thereby facilitating downstream analyses. The error correction module effectively rectifies translocation, inversion, and debris errors at their identified locations. Subsequently, based on the model’s detection of the number of chromosomes, the AutoHiC split module divides the previously corrected genome results into the final scaffold genome at the chromosome level. In the final stage, a visual report is generated, showcasing the genome before and after error correction. This report plays a pivotal role as it enables users to assess genome quality, examine pertinent indicators, and review details of the error correction process for subsequent analyses.

### Assembly report

The detailed assembly result report is the bridge between the user and the assembly result. It allows users to fully understand their own data. In addition, the comprehensive assessment of AutoHiC corrections is facilitated through a detailed result report, serving as a pivotal tool for validation. This report encompasses essential aspects of the final genome assembly, error rectification intricacies, and pertinent details. Structurally, the report is compartmentalized into four sections, each strategically addressing distinct facets of the genomic correction process.

The initial segment of the report furnishes fundamental genome attributes, encompassing pivotal parameters such as genome size, N50 value, L50 value, Hi-C anchor rate, scaffold count, and GC content. This segment is instrumental in offering an overview of the genome’s characteristics post AutoHiC correction, serving as a foundation for subsequent analysis and interpretation (Supplementary Figure 1).

The subsequent section of the report undertakes a comparative evaluation of the genome-wide interaction heatmaps before and after the error correction process. This visual juxtaposition provides clear insight into the efficacy of AutoHiC corrections in enhancing the precision and reliability of genomic interactions, bolstering the overall integrity of the assembly (Supplementary Figure 2).

The third section delves into the granular intricacies of the error correction process. It meticulously outlines the specifics of identified errors, encompassing their dimensions, positional coordinates within the genome, and the visual representation of the nature and context of the errors. This detailed exposition not only facilitates an in-depth comprehension of the corrective process but also supports downstream investigations and potential optimizations (Supplementary Figure 3). For example, when studying a 3D genome, it is possible to focus on whether these regions have an impact on the formation of the 3D structure of the genome.

Finally, the report culminates with an appendix containing supplementary information. This section is instrumental in recording and tracking the dynamic evolution of errors during the correction process. It encapsulates the fluctuation in error count and the consequential impact on chromosome length determinations. This comprehensive archive of data affords deeper insight into the iterative nature of AutoHiC corrections, enriching the overall picture of the algorithm’s performance (Supplementary Figure 4). In essence, the results report stands as a cornerstone for the comprehensive assessment of the efficacy of AutoHiC in refining genome assemblies. Through its structured segments, AutoHiC provides a multifaceted lens to scrutinize genomic attributes, error rectification outcomes, and their subsequent implications, thereby underscoring the robustness and applicability of AutoHiC in contemporary genomics research.

### The AutoHiC model

Accurately identifying the type of error and where it occurred is key to correcting assembly errors. We developed the AutoHiC tool, which can eliminate the effects of complex features caused by assembly errors, including error size, resolution, and color gamut (Fig. 1b). AutoHiC is a deep neural network that utilizes two-stage object detectors to enhance the detection of genome assembly errors and extract error features by leveraging Hi-C data (Methods). AutoHiC is based on the Swin Transformer^37^ architecture, which incorporates self-attention mechanisms. We chose this architecture as a starting point based on substantial evidence indicating that quantized architectures presently yield superior image representations.

The AutoHiC model employs a streaming sliding window approach (Methods) to scan the entire Hi-C contact map. The contact map is then converted into an image to capture error signals, and corresponding log files are generated for subsequent error processing. A trained neural network (Methods) is used to extract error features for each image, encoding the location, type, and score of the error in the image. High-confidence error predictions are refined and mapped back from the image to the scaffold genome coordinates, and this information is utilized in the error correction module (Methods).

### Principles of AutoHiC’s algorithm

We subsequently assessed the capability of the AutoHiC algorithm in identifying critical features such as translocations, inversions, debris, and chromosomes, which are pivotal for studying and rectifying errors in genome assembly. The AutoHiC model, as depicted in Figure 1b, employs a two-stage object detection framework within a deep neural network. It effectively pinpoints and locates erroneous regions in misassembled scaffolds, subsequently rectifying them using information provided by the model in conjunction with the Hi-C interaction matrix. AutoHiC retrieves pertinent interaction matrix data based on error position information provided by the model and employs the peak algorithm to precisely identify the erroneous insertion position and boundary.

For translocation errors, the algorithm extracts the regions characterized by the error detection model. Interaction curves are then generated for the corresponding regions. It can be clearly seen that the interaction law of the interaction curve corresponds to the interaction heatmap and can indicate the location where the translocation error occurs and the site that needs to be inserted (Fig. 2a). Due to the presence of interfering signals, AutoHiC eliminates peaks at its own site and filters redundant peaks to determine the exact insertion site of the translocation error. In addition, AutoHiC shifts the sequence of the region where translocation occurs to the corresponding insertion site. As seen from the interaction heatmap and the interaction curve (Fig. 2b), after the AutoHiC adjustment, the interaction is more in line with the interaction law, the translocation error is also eliminated on the heatmap, and the interaction curve returns to normal. Inversion errors have their own specific characteristics in both interaction heatmaps and interaction curves (Fig. 2c). On the interaction heatmap, the area where the inversion error occurs is shaped like a butterfly, and the corresponding interaction curve has a peak area of a certain length. AutoHiC calculates the length of the inversion error based on the area of the peak on the interaction curve and then adjusts the sequence in that area in the opposite direction. After AutoHiC adjustment, the features on the interaction heatmap and the peaks on the interaction curve are eliminated, indicating that the algorithm can adjust the inversion error well (Fig. 2d). Debris errors are usually short in length and appear as blanks on the interaction heatmap and as a region of zero interaction on the interaction curve (Fig. 2e). Similar to the inversion error, the algorithm calculates the error length based on the interaction curve and then deletes this sequence from the genome. Compared to the interaction heatmap and interaction curve before adjustment, the blank area on the interaction heatmap after adjustment is eliminated, and there is no segment with an interaction value of 0 on the interaction curve (Fig. 2f).

**Fig. 2:**
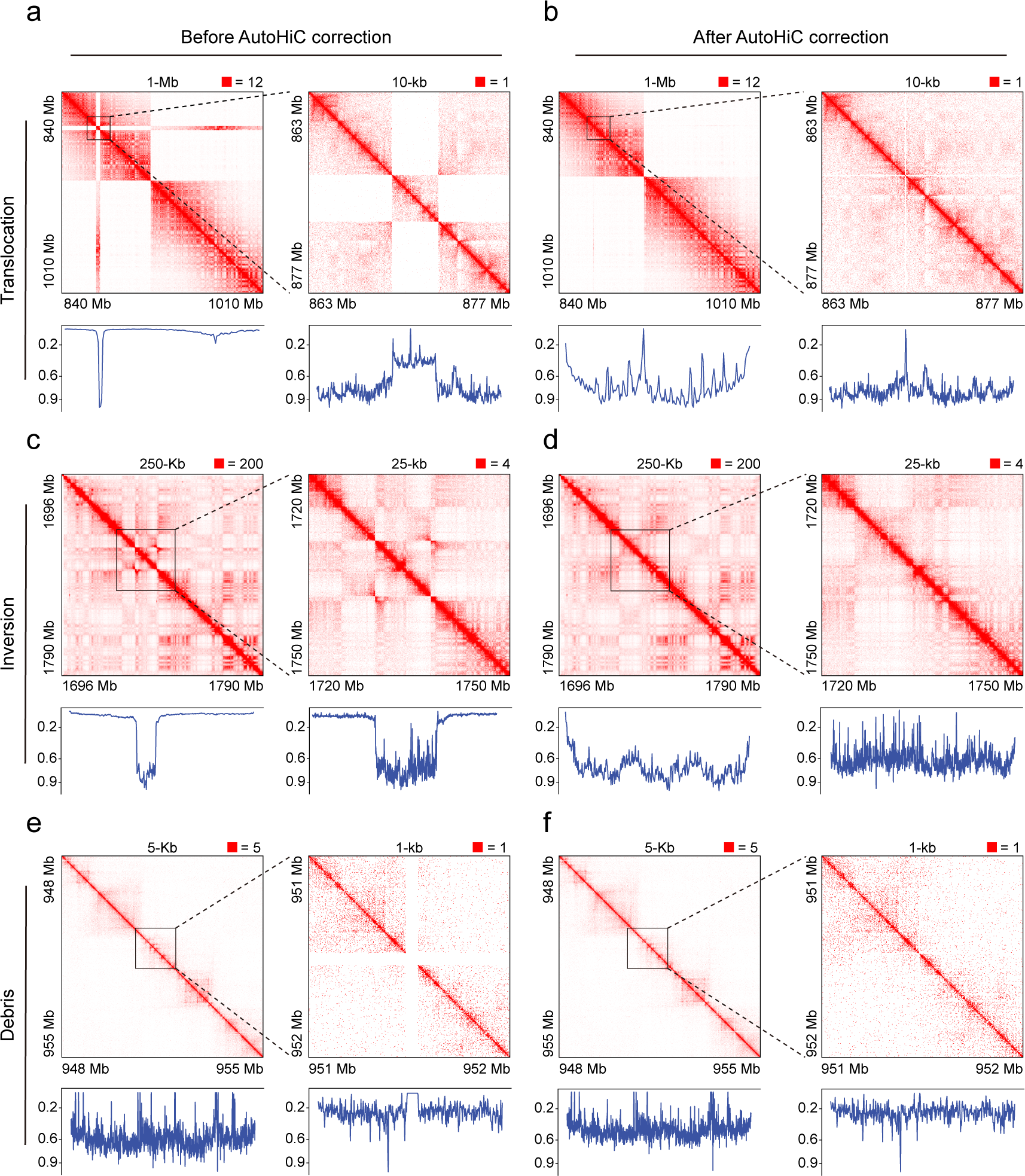
Comparison of AutoHiC error correction effects. **a, c, e** Interaction heatmaps and interaction curves with assembly errors. The interaction heatmap is at two resolutions, with the high resolution on the right. The resolution and interaction boundaries are marked at the top. The lower boundary has a marker indicating the extent of the interaction area. Interaction curves are generated from the interaction matrix used by the interaction heatmap. Interaction values have been normalized to 0-1. **b, d, f** Interaction heatmaps and curves after error corrections without assembly errors.

In addition to identifying and rectifying genome misassembly, AutoHiC can also be applied to chromosome splitting (Supplementary Figure 5). Upon correcting all errors, AutoHiC infers the number of chromosomes in the genome by utilizing the global interaction heatmap and subsequently assigns the genome sequence to their respective chromosomes (Methods).

The exceptional capability of AutoHiC in accurately identifying and positioning both large and small translocations, inversions, debris, and chromosomes underscores its high potential for studies involving genome structure comparison. Furthermore, its proficiency in detecting and correctly assigning error regions with low rates of false positives sets AutoHiC apart from previous methodologies.

### Performance evaluation of AutoHiC

We present the training results of AutoHiC for assembly error detection and chromosome detection (Supplementary Figure 6). The accuracy and loss during model training are depicted to illustrate the convergence and fitness of the model to the training data after 200 epochs (Fig. 3a, b), indicating an effective learning process. To assess the model’s performance comprehensively, we employ the confusion matrix and precision–recall curve. The confusion matrix provides valuable insights into the model’s performance across different classes (translocation, inversion and debris). Evaluating the model’s performance for each specific class offers an internal perspective on its effectiveness, yielding a more nuanced assessment compared to overall accuracy. Notably, the confusion matrix (Fig. 3c) indicates that the model predicts translocation, inversion, and debris errors with image accuracy exceeding 90%, while inversion prediction achieves 85% accuracy. The instances where the model incorrectly predicted images predominantly occurred in complex situations. The consistent patterns observed in the confusion matrix align with the dataset’s regularity, validating the reliability of the AutoHiC model for error detection.

**Fig. 3:**
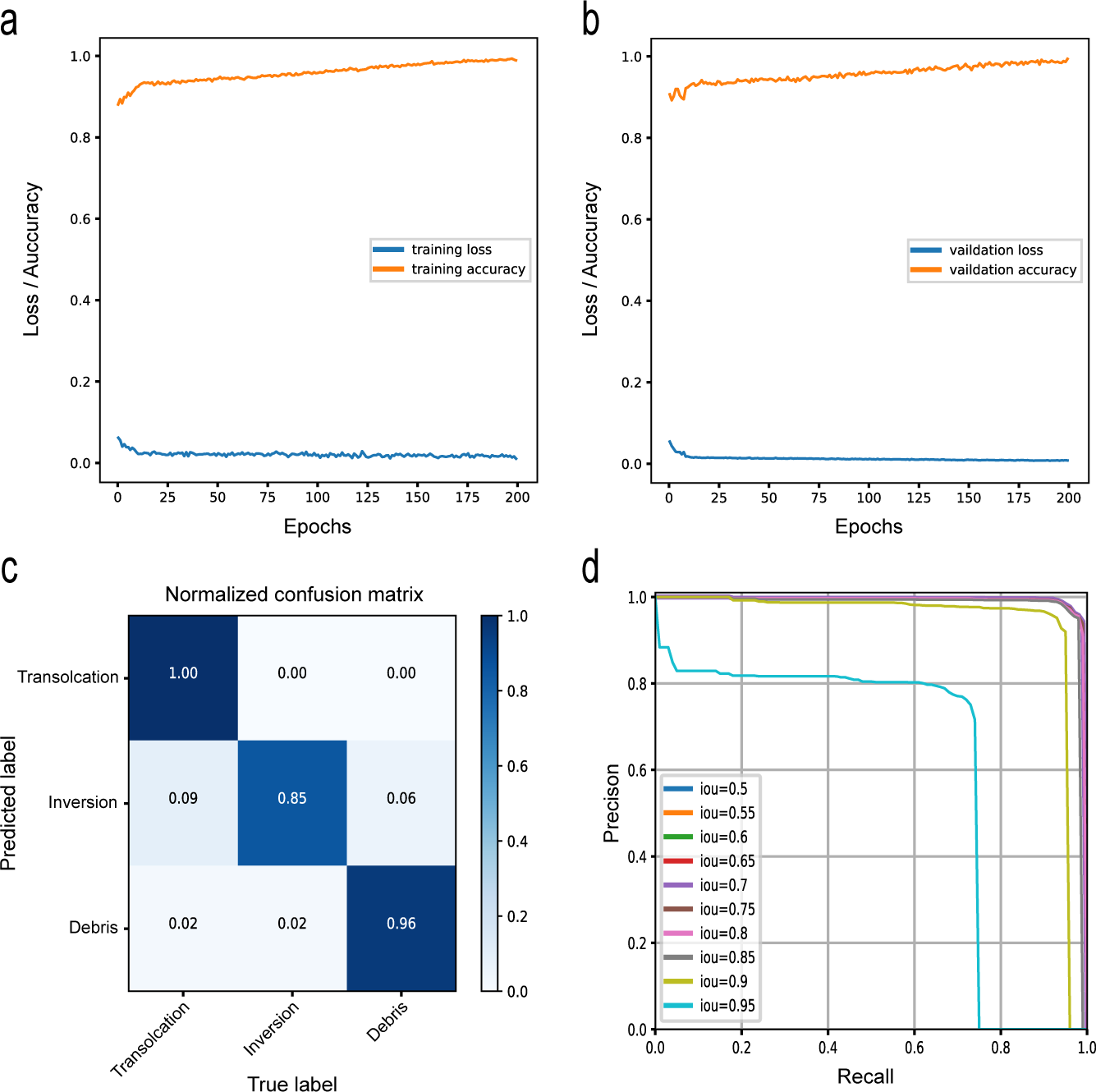
AutoHiC model effect verification. **a, b** Changes in model accuracy and loss rate during training and validation. On the left is the training process and on the right is the verification process. The lower curve is the training loss and the upper curve is the training accuracy. **c** Confusion matrix results for the AutoHiC error detection model. The horizontal axis of the graph represents the actual error, and the vertical axis represents the model-predicted error. The diagonal line is the correct prediction result, and the darker the color of the cube, the more images are correctly predicted. **d** Precision recall (PR) curve for the different IOU thresholds as in panel. Curves without color represent PR results at different IOU thresholds.

Furthermore, we compared the performance of the AutoHiC model using the precision-recall curve (Fig. 3d) and the area under the curve (AUC). Multiple precision-recall pairs were calculated by varying thresholds, and their visualization allowed us to compute the AUC using the composite trapezoidal rule. A higher AUC value, ranging from 0 to 1, indicates superior model performance. The precision-recall curve demonstrates that when the threshold is set between 0.5 and 0.95, the area under the curve approaches 1, highlighting the model’s strong performance.

### AutoHiC outperforms in improving assembled genome quality

To further benchmark^38^ the performance of AutoHiC, we conducted a comparison of AutoHiC with several competitive and representative tools, including 3D-DNA, SALSA2, YaSH, and Pin_hic (Supplementary Note 1; Supplementary Table 1). Our primary goal was to evaluate the effect of AutoHiC on improving genome continuity. To achieve this, we selected five species: *Caenorhabditis elegans*, *Arabidopsis thaliana*, *Drosophila melanogaster*, *Danio rerio*, and *Homo sapiens*. The selection of these species is based on the fact that they represent a good representation of currently studied model species, including plants, animals, and humans. Currently used for continuity assessment is N50^39^, which indicates the degree of continuity of the genome assembly, which is defined by the length of the shortest contig for which longer and equal length contigs cover at least 50 *%* of the assembly. Higher N50 values indicate better continuity. Additionally, the L50 value corresponds to the N50 value and indicates the number of contigs (or scaffolds) required to achieve the N50 value. Lower L50 values indicate better continuity. Therefore, we calculated the Nx (Fig. 4a, b, c, d, e; Supplementary Table 2) and L50 (Supplementary Figure 7) of these software assembly results using QUAST (Methods). Unexpectedly, we observed from the results that the N50 values of YaSH and Pin_hic were very large, almost approaching the size of the entire genome, which is clearly problematic (Supplementary Note 2; Supplementary Table 3). Consequently, we analyzed the assembly results obtained by these two software programs separately (Supplementary Figure 8; Supplementary Table 4) and found that they merged the sequence of the entire genome into a single chromosome. After excluding these problematic outcomes, AutoHiC performed remarkably well on the five test species, exhibiting the highest N50 value. Compared to the raw data and the output of other software programs, AutoHiC improved the continuity by approximately 18-fold (compared to Contig) and 7-fold (compared to SALSA2).

**Fig. 4:**
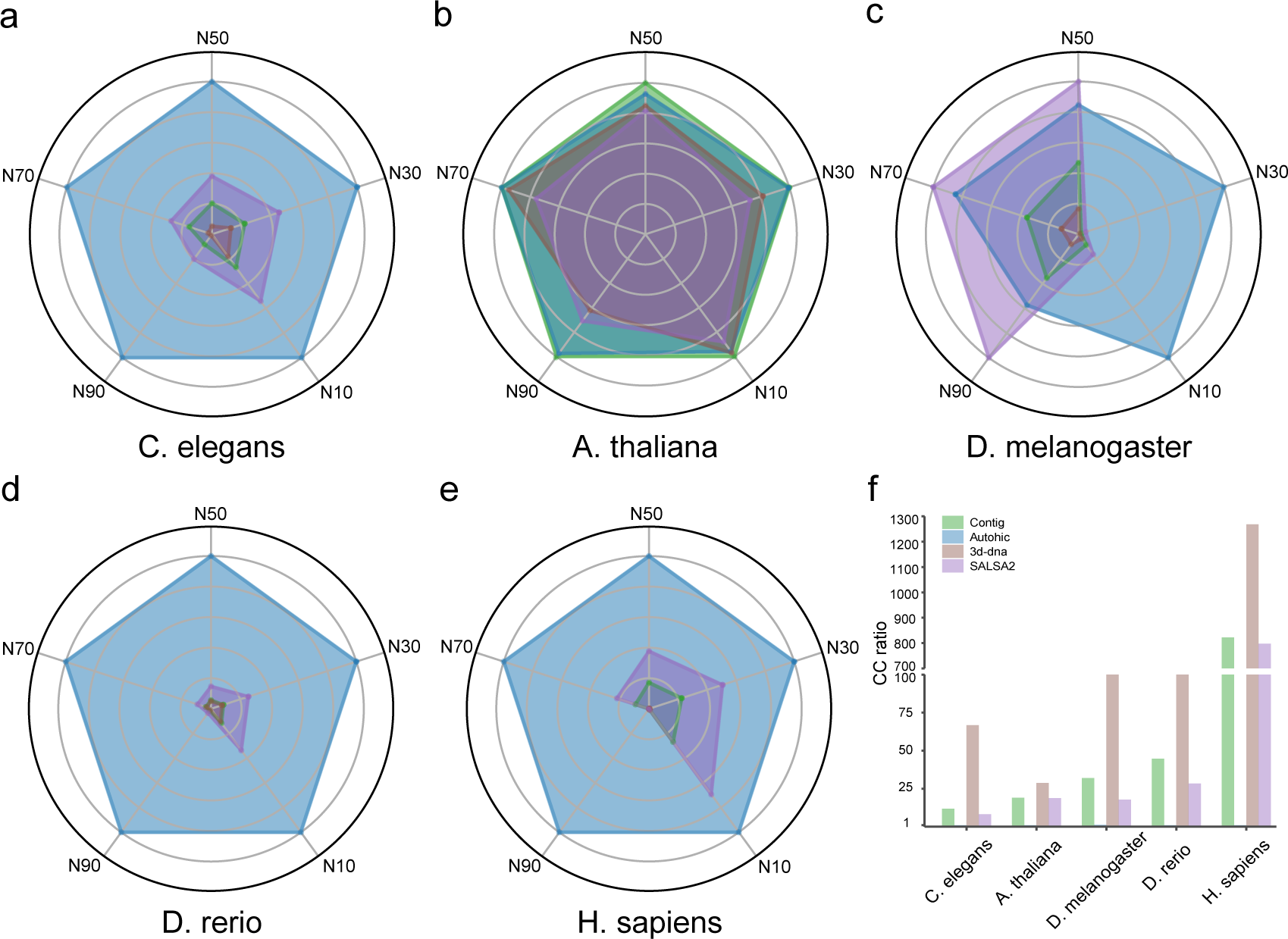
Comparison of genome continuity benchmarks. **a, b, c, d, e** The radar chart shows the continuity of the assembly results. Different colors represent different software programs. The radar chart shows N10, N30, N50, N70 and N90. Different species are shown separately. **f** Histogram of CC rate values of different software contig results. It contains the results of the contig, and different colors represent different software.

For a more robust assessment of continuity, we also introduce the CC ratio^40^, a score that intuitively reflects continuity regardless of contig length or intrinsic chromosomal length. The results (Fig. 4f) showed that the output from all the software tools had significantly higher CC values, with the exception of AutoHiC, which remained at 1. This shows that the genome assembled with AutoHiC had high continuity and was at the chromosome level. Based on the aforementioned evaluation results, AutoHiC demonstrates superior performance compared to the other four scaffolds in terms of enhancing genome continuity.

### Validation of AutoHiC results using the T2T genome

We selected five T2T genomes^41,42^ (*Caenorhabditis elegans*, *Arabidopsis thaliana, Bombyx mori, Oryza sativa* and *Homo sapiens*) to evaluate the effect of AutoHiC on improving genome continuity and accuracy (Methods). The selection of the T2T genome is mainly based on the following rationale. First, the T2T genome, as the gold standard, can truly reflect the actual situation of the genome and can then be compared with the adjusted results from AutoHiC. Second, we can check whether AutoHiC is overtuned by comparing the number of structural variations before and after AutoHiC correction. Finally, this approach also enables validation of whether AutoHiC-adjusted genomes are closer in accuracy to T2T genomes than genomes from conventional methods.

Initially, we utilized QUAST to analyze the assembled genome before and after AutoHiC error correction (Supplementary Table 5). Remarkably, AutoHiC significantly improved genome continuity (Fig. 5a, b; Supplementary Figure 14), with the number of scaffolds in the corrected genome approaching the number of chromosomes (approximately equal to the number of chromosomes). The NGA50 and NG50 values were also greatly improved. Additionally, we employed MUM&Co to compare the assembly results with the T2T reference genome and quantified the number of structural variations (Fig. 5c; Supplementary Figure 14). This evaluation partly reflects the performance of the assembly software in terms of accuracy and completeness. Notably, compared to the assembly results from other software, the AutoHiC assembly results showed the lowest number of structural variations, highlighting the superiority of AutoHiC in improving genome accuracy and indicating that there was no overtuning of AutoHiC. If AutoHiC is overtuned, there will be a large difference between the genome assembly results of AutoHiC and the T2T genome, and more structural variations will be found after the whole genome comparison between them.

**Fig. 5:**
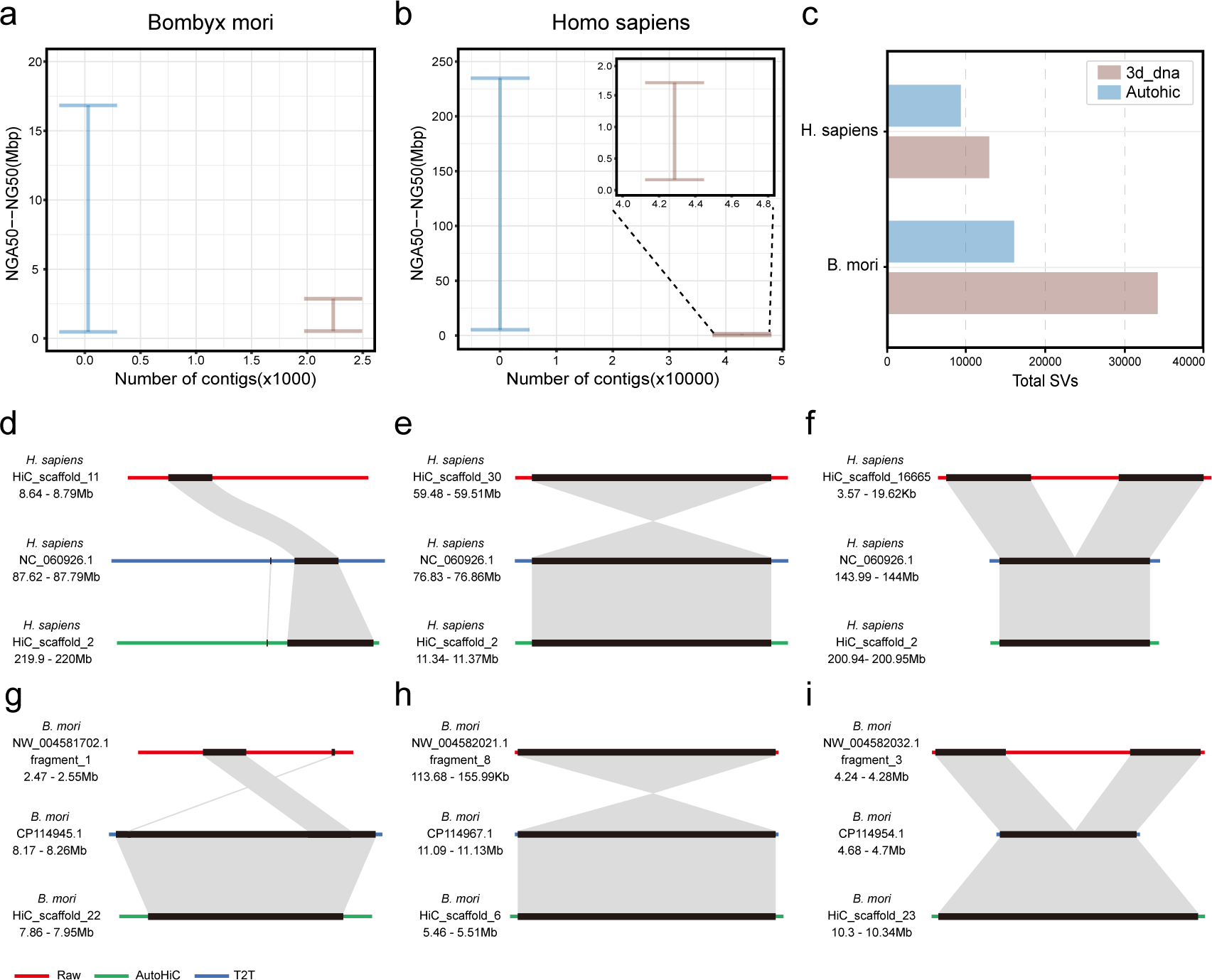
Assessing the performance of correcting the T2T genome. **a, b** Comparison of the contiguity of assemblies scaffolded by 3D-DNA and AutoHiC as measured by QUAST. The Y-axes show the range of NGA50 to NG50 lengths to indicate the uncertainty caused by true genomic variation between the individual and the reference genome. **c** Comparison of the accuracy of assemblies scaffolded by 3D-DNA and AutoHiC as measured by MUM&Co. The X-axis shows the total number of structural variations to show AutoHiC’s ability to correct errors. **d, e, f, g, h, i** Collinearity block plots before and after three error corrections for translocation, inversion and debris. Above is the uncorrected collinearity with the T2T genome. Below is the AutoHiC-corrected collinearity with the T2T genome in the same region. The names and positions of the sequences are marked on the left.

In addition, to verify the accuracy of the AutoHiC error correction and the degree of consistency between the alignment results and the T2T genome, we compared the *Homo sapiens* and *Bombyx mori* genomes with the T2T reference genome by synteny analysis (Methods). Based on the structural error information detected by MUM&Co, we selected three types of error corresponding sequences and constructed a block collinearity map. For translocation errors (Fig. 5d, g), it can be seen from the collinearity map that, using the T2T genome as a reference, the region where the translocation error occurred was where the sequence had obvious placement errors. However, after AutoHiC adjustment, this issue was eliminated, the position of the sequence returned to normal, and there was a better collinearity result in the same region. For inversion errors (Fig. 5e, h), the regions where the translocation errors occurred showed opposite sequence characteristics on the collinearity map, and after correction by AutoHiC, the collinearity with the T2T genome was more consistent. Additionally, there was a debris error (Fig. 5f, i). From the collinearity map we can see that there is a fragment without collinearity in the original genome, resulting from accidental insertion or other methods, but it is certain that it does not belong to this region. Therefore, AutoHiC would adjust the position of this fragment so that the genome has a higher identity with the T2T genome.

In conclusion, AutoHiC significantly improves genome assembly accuracy. We anticipate that when the AutoHiC model is integrated with genome assembly software, it will more effectively address Hi-C assembly errors.

### Extending AutoHiC to more complex situations

To further verify the performance of AutoHiC and demonstrate its broad applicability and robustness, we applied AutoHiC to seven more extensive datasets from DNA Zoo and NCBI: very large genome (*Schistocerca americana*), large number of chromosomes (*Chiloscyllium punctatum*), plants (*Oryza sativa*), polyploidy (*Arachis hypogaea*), etc.

To assess genome continuity, we calculated the N50 value, L50 value and CC ratio (Supplementary Figures 15, 16, 17) and observed a significant improvement in genome continuity after AutoHiC correction. Moreover, we developed an independent error rate index (Methods) to facilitate the visualization of the correction effect. Typically, translocation and inversion errors disappear after one round of correction, while some debris errors remain (Supplementary Figure 18). Some specific genomes required four corrections to achieve error elimination. The error rate is computed based on the error length relative to the genome size. As a result, its value exhibits minimal fluctuation; however, a discernible downward trend is evident.

To compare the changes before and after processing, the Hi-C interaction heatmaps of the seven species are presented before and after correction (Figure 6a, b, c, d, e, f). It is evident that the assembly errors observed in the Hi-C interaction heatmap prior to correction were eliminated, and redundant sequences were removed. In addition, some interaction heatmaps were visually compared before and after error correction to clearly illustrate the changes (Supplementary Figures 19, 20, 21).

**Fig. 6:**
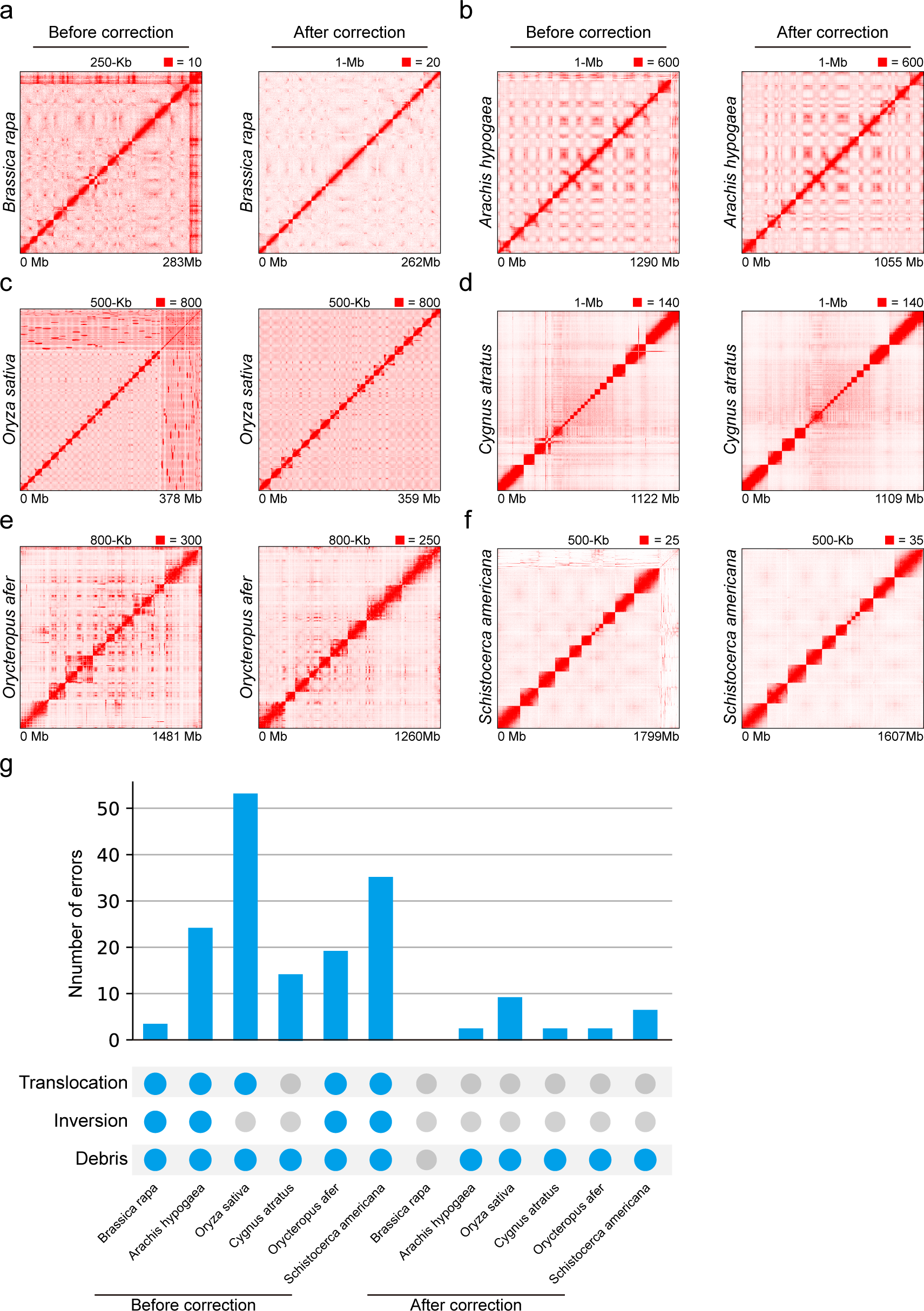
Genomic dataset extension validation. **a, b, c, d, e, f** Global interaction heatmaps before and after AutoHiC error correction. The left is before correction. The right is corrected. Resolution and interaction thresholds are marked at the top. The lower value is the genome length (ratio). **g** Histogram of the number of errors before and after AutoHiC error correction. The histogram above is the sum of the three error numbers. A colored circle below indicates the presence of such errors. The 6 on the left are before AutoHiC corrects the error, and the ones on the right are after the correction.

The changes in the number of assembly errors were recorded during the correction of the genome with AutoHiC. After error correction by AutoHiC, the number of errors in the Hi-C heatmap was significantly reduced. Due to debris errors in the Hi-C heatmap, no unified conclusion was reached. Consequently, AutoHiC did not entirely delete this part of the sequence. Users can choose to clear all errors based on their needs.

The above results clearly demonstrate that the correction of genomic misassemblies by AutoHiC significantly improves both continuity and correctness in terms of genome assembly. In summary, we demonstrate the effectiveness of AutoHiC when applied to complex genomic datasets and highlight its broad applicability, which is critical for downstream analyses.

## Discussion

In this study, we present an innovative deep learning-based tool called AutoHiC, which is designed to identify and correct misassembled contigs within assemblies. We highlight its profound impact on enhancing both genome continuity and accuracy across a diverse array of datasets characterized by varying complexities. AutoHiC stands apart from prior methodologies in two main pivotal aspects. First, unlike conventional assembly approaches^43–46^ that predominantly rely on 1D or 2D data from Hi-C, AutoHiC harnesses the potency of higher-dimensional Hi-C information to facilitate error correction during the genome assembly process. The traditional assembly method uses only the relationship between the interaction matrix and the genome sequence for assembly. On this basis, AutoHiC converts the two-dimensional interaction matrix into a three-dimensional interaction heatmap to detect and correct assembly errors. A genome can be assembled more comprehensively and accurately, compensating for certain shortcomings of the traditional method. Second, AutoHiC assembly pipeline operates in a fully automated manner (depicted in Figure 1a); neither error correction nor chromosome splitting (as illustrated in Figure 2 and Supplementary Figure 5) necessitates manual intervention—only the provision of genome and Hi-C data. Furthermore, AutoHiC provides an explicit report for each assembled genome, a feature with substantial potential value for downstream analyses. Notably, owing to its algorithm agnostic nature, AutoHiC exhibits versatility in accommodating varying sizes and styles of genomes, transcending species-specific constraints (as evidenced in Figure 6). Leveraging the computational resources optimally, AutoHiC capitalizes on the multithreading capabilities of both central processing units and graphics processing units, thereby delivering swift outcomes.

There are several directions that hold promise for further improvements to AutoHiC. First, AutoHiC takes into account the rapid iteration of the scaffolding algorithm at the beginning of the design so that AutoHiC can seamlessly change the scaffolding to take full advantage of the scaffolding. The combination of AutoHiC and scaffolding algorithms is therefore a promising direction for future research, leading to effective approaches for reconstructing genomes from sequencing data with higher quality and completeness. In fact, interaction types have a wide variety of forms and appearances, reflecting a variety of states. This rich diversity is much more difficult to model and analyze than, for example, sequence variability. Therefore, it may be necessary in the future to increase the type of error detection to correct assembly errors more comprehensively.

Anticipating future improvements, AutoHiC strategically accounts for the rapid iteration of scaffolding algorithms in its design, enabling seamless alignment with evolving scaffolding techniques. This convergence of AutoHiC and scaffolding algorithms offers a promising trajectory for subsequent research, leading to advanced strategies in genome reconstruction with heightened quality and comprehensiveness. Importantly, the diverse array of interaction types, with their intricate forms and manifestations, present a more challenging modeling and analytical scenario than sequence variability presents. Consequently, augmenting the spectrum of error detection to encompass a wider array of types may be warranted for more comprehensive assembly error correction.

With the escalating availability of high-quality Hi-C datasets, we envision that the potency of AutoHiC will continue to flourish. To encapsulate, AutoHiC serves as a tool, showcasing the transformative potential of the transformer architecture in modeling biological sequencing data. As the landscape of genome project burgeons and achievements in sequencing technology scale to new heights, we assert that AutoHiC is primed to make a substantial contribution to the genomics domain, liberating scientists from labor-intensive manual curation.

## Methods

### AutoHiC workflow

AutoHiC comprises three primary steps. To initiate the process, users are required to prepare preexisting contig data, Hi-C sequencing data, and details pertaining to the restriction enzymes employed in the Hi-C experimental setup. The initial stage involves the invocation of Juicer to perform the alignment of Hi-C reads onto the contig genome, thereby securing a robust alignment outcome. Subsequently, an intermediary file is generated, encapsulating a comprehensive spectrum of genome-wide interaction information. This intermediary file, serving as a foundation, is then seamlessly integrated into the 3D-DNA platform to orchestrate a refined assembly of contigs into scaffold structures. Notably, this phase engenders an array of outputs, among which the hic, assembly, and scaffolds files hold pivotal significance as input materials for AutoHiC’s subsequent error correction module.

The second phase ushers in AutoHiC’s error identification and rectification module, a cornerstone of its innovative workflow. By leveraging the preliminary outputs generated thus far, this module detects assembly anomalies, iteratively navigating through the error correction process. This error correction endeavor is complemented by the allocation of sequences to their respective chromosomal contexts, determined through predictive analytics concerning chromosome count.

The culmination of the entire assembly process resides in the third and final stages. Here, a holistic synthesis of all relevant assembly information materializes, culminating in the fruition of the adjusted genome. Finally, the genome is statistically analyzed and a comprehensive results report is generated.

A notable caveat pertains to the prevalent utilization of 3D-DNA as a linchpin in the initial phase of AutoHiC’s automated assembly process. This selection is primarily rooted in the software’s preeminence and pervasive adoption within the domain of genome assembly. However, it is imperative to underscore that for users operating distinct assembly software, the interaction outcomes can be transformed into requisite formats, thereby enabling the independent utilization of AutoHiC for assembly correction and associated tasks.

### Error identification

AutoHiC employs a specific methodology known as the Hi-C contact matrix to accurately pinpoint instances of misassembly breakpoints within the genome. The construction of the Hi-C contact matrix involves the alignment of contact reads against the reference genome, yielding invaluable insights into both intra- and interchromosomal interactions. These assumptions are rooted in the empirical observation that Hi-C interaction maps faithfully depict the actual physical distances separating different genome sequences. In essence, AutoHiC transforms the Hi-C interaction map into multichannel images, adeptly capturing a diverse array of error-indicative signals. Subsequently, a network of transformers is harnessed to predict the precise error type and genomic locus depicted in each image. Through this intricate process, AutoHiC effectively identifies instances of misassembled contigs.

In the operational context, AutoHiC converts the interaction heatmap into a three-channel RGB image through a sliding window approach tailored to the resolution and dimensions of the interaction matrix. Each image, enriched with interaction-related information, is then fed into a pretrained error detection model. This model scrutinizes the presence of assembly errors within the image and, if detected, records pertinent information about the type and location of the error. AutoHiC integrates an inherent information conversion script that translates error positional data identified within the image into corresponding genomic coordinates. This information is subsequently utilized by the AutoHiC error correction module to effectively rectify the errors.

### Error correction

The AutoHiC algorithm’s pivotal component is its error correction module. Operating as the algorithm’s core, this module is designed to rectify errors within the genome by leveraging information about their specific locations. Functioning as a subsequent step following error detection, it is primarily responsible for addressing three major categories of errors: translocation, inversion, and debris. Among these, translocation errors present the most intricate challenge. Their resolution entails extracting the corresponding interaction matrix within the identified interval and subsequently transforming this matrix into a peak map. Through the application of the peak algorithm, the insertion interval for the translocation error is meticulously identified. Ultimately, the pertinent genomic sequence undergoes precise adjustments to restore accuracy.

In contrast, inversion and debris errors do not necessitate the same level of complexity in terms of locating insertion points. Rather, their rectification primarily involves refining the sequences associated with the genomic region. In instances where scaffold assemblies exhibit high-quality attributes, a single iteration of AutoHiC correction often proves sufficient to comprehensively rectify all identified errors. This iterative strategy contributes significantly to heightening the algorithm’s accuracy and overall reliability.

The meticulous orchestration of the error correction module within AutoHiC underscores its critical role in enhancing the fidelity of genome assemblies. By systematically addressing various types of misassemblies, AutoHiC ensures the integrity of genomic data, which is imperative for downstream analyses and the advancement of genomics research.

### Peak algorithm

The main task of the peak algorithm is to find insertion sites of translocation errors. It extracts the interaction matrix

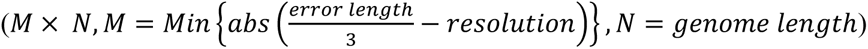

with the best resolution based on the incorrect length.

Then, the peaks in the interaction matrix are extracted, and after excluding the position of the error and the peaks (*Max*{*i*_1_, *i*_2_, *i*_3_, … *i*_*n*_}) of the accessory, the site selected according to the maximum value is the insertion site of the translocation error.

### Chromosome number prediction

Chromosome number prediction relies on the global interaction heatmap (Supplementary Figure 5). Initially, the error-corrected interaction matrix, which has been purged of redundant sequences, is transformed into a visual representation through a visualization tool. The chromosome detection model then analyzes these generated visual representations to ascertain both the count and size of chromosomes. Subsequently, AutoHiC allocates the genomic sequences to their respective chromosomes according to the deduced information.

### Model architecture

AutoHiC consists of four essential components: backbone, neck, densehead, and roihead. To enhance the extraction of error features from images, we employ the Swin Transformer as the backbone network of AutoHiC. This network adopts a layered transformer structure, dividing the image into different blocks and performing self-attention operations within each block. Such a hierarchical architecture enables efficient handling of large-scale Hi-C interaction images. To mitigate computational complexity, the model incorporates a windowed attention mechanism, confining self-attention operations within local windows of each block rather than performing global computations. Additionally, a deep cross-local area attention mechanism is introduced to establish long-range dependencies among different blocks, capturing a broader range of contextual information.

To facilitate feature transformation and fusion, AutoHiC integrates the feature pyramid network^47^ (FPN) as a neck. FPN is adept at handling visual tasks at various scales and layers, making it an ideal choice for AutoHiC architecture. Finally, for the precise screening and localization of extracted error features, the Cascade Region of Interest (RoI) Head and Region Proposal Network^48^ (RPN) Head are employed. For comprehensive technical details, readers can refer to the implementation code.

### Streaming sliding window

The sliding window algorithm is based on the set size to extract the local matrix by sampling the interaction matrix extracted at different resolutions. For different resolutions, the size of the sliding window is *M* (*resolution* ∗ 700), and the sliding step is *N* (*resolution* ∗ 400). The whole process is performed along the diagonal of the interaction matrix from top left to bottom right.

### Datasets

The dataset used in this article is divided into two parts. One part is the training data of the model, mainly the original interaction information in hic format. Another part is test data used to evaluate the real data of AutoHiC performance and application range, which includes genome, Hi-C sequencing data, etc. All the data utilized in the study presented in this article are sourced from DNA Zoo and NCBI. Details regarding the genome and Hi-C sequencing data can be found in Supplementary Table 17. The raw data used for model training are obtained from DNA Zoo, and specific information is available in Supplementary Table 18.

### Generation of training data

The data used for training the model are primarily generated and simulated based on collected Hi-C data. The generated data are created using a script that follows predetermined length gradients and resolutions, while the simulated data are produced by randomly adjusting interaction matrix values derived from actual data, resulting in data that closely resemble real-world observations. The color thresholds applied to heatmaps are determined by extracting the 95th quantile of the interaction matrix distribution.

### Image data preprocessing

The initial data are subjected to preprocessing to convert them into localized interaction heatmaps represented in jpg format. Due to the substantial volume of generated data, direct utilization for model training becomes unfeasible. To address this, a classification network is implemented to curate usable images from the generated dataset. Subsequently, manual annotation was performed on the selected data using the labelme (v.5.3.1) tool. Prior to the training phase, data augmentation procedures are applied, encompassing transformations such as flipping, rotation, scaling, cropping, and shifting. These augmentations are designed to amplify the diversity and robustness of the training dataset.

### Classification network

The training data for the error detection model come from the screening of the classification model. The classification model is used for preliminary screening of generated and simulated Hi-C data for subsequent manual screening and labeling. The classification model is based on EfficientNetV2^49^. Its training data come from errors that were originally detected by humans.

### Model training

The dataset is partitioned into two distinct subsets, with approximately two-thirds of the data serving as the training set to iteratively refine the model parameters. The remaining one-third is allocated as the test set, a set-aside corpus intended to meticulously assess the performance of the trained models. It is crucial to underscore that the test set remains entirely unseen and untapped during the training regimen, thus guaranteeing an impartial appraisal of the model’s efficacy.

### Model evaluation

Furthermore, we ascertained the generalization proficiency of AutoHiC through an evaluation conducted on an independent test subset that remained entirely segregated from the training process. In appraising the detector’s efficacy posttraining, we leverage both the confusion matrix and the area under the precision-recall curve (AUPRC). For each pair of adjacent precision and recall rates, treat them as the upper and lower bases of a trapezoid and calculate the area of the trapezoid to estimate the area under the PR curve at the current stage. For all trapezoids, calculate their areas and add them together to obtain an approximation of the area under the PR curve. During the cross-validation procedure, we judiciously select hyperparameters that yield the highest AUPRC, thus ensuring the optimal configuration settings.

### Algorithm verification

To assess the feasibility of the AutoHiC error correction algorithm, we employed a comparative approach involving the generation of interaction heatmaps and interaction curves before and after the error correction process. The interaction heatmap was produced using pyStraw, targeting two distinct resolutions corresponding to the intervals where errors occurred. Notably, the intervals remained consistent between the pre- and postcorrection phases. Higher resolutions were adopted to amplify the precision of erroneous details. Simultaneously, the interaction curve was derived from the interaction matrix, specifically by focusing on the positions where errors were identified. While the data within the interaction matrix remained unnormalized, the extracted interaction matrix underwent normalization to achieve a range of 0 to 1, subsequently facilitating the generation of the associated interaction curve.

### Benchmarking the performance of AutoHiC

All the compared software programs were obtained, installed, and evaluated following the procedures outlined in the provided publication sources. When assessing the performance of different software programs on the same species (*Caenorhabditis elegans*, *Arabidopsis thaliana*, *Drosophila melanogaster*, *Danio rerio*, and *Homo sapiens*), uniform datasets were employed to ensure consistency. The input data for software such as SALSA2, YaSH, and Pin_hic consisted of BAM or BED files generated by the HiC-Pro^50^ pipeline. Parameter configurations for each of the compared software programs were maintained at their default settings as stipulated by the original documentation for each program.

The genome continuity assessment predominantly relied on key metrics such as the N10-N90, L50, NGA50, and NG50 values and the CC ratio. Specifically, the N10-N90 and L50 metrics were computed utilizing the QUAST software package (v5.2.0). The parameter settings for all compared software were retained in their default states, as prescribed by the original documentation. The CC ratio, representing the proportion of contig count to chromosome count, was established manually.

For the quantitative analysis of structural variations, an appropriate reference genome was selected from Supplementary Table 12. The analysis was conducted using MUM&Co software (v3.8), maintaining parameter configurations in line with the default specifications as provided in the original documentation.

### Synteny analysis

For comparative analysis, we assessed the genomes assembled by 3D-DNA and those corrected by AutoHiC against their respective T2T reference genomes. This evaluation involved utilizing the MUM&Co tool to capture structural variation information. Leveraging the structural variation data from MUM&Co, we employed NGenomeSyn^51^ (v.1.41) to visualize structural variation regions within the 3D-DNA assembly outcomes. Subsequently, we harnessed SeqKit^52^ (v.2.5.1) to extract sequences containing structural errors, followed by employing blastn to conduct sequence comparisons between these sequences and the genomes adjusted by AutoHiC. This comparison enabled the derivation of corresponding sequence positional information. Criteria for filtering the comparison results were primarily guided by sequence length and alignment consistency. Ultimately, we utilized SeqKit to extract the aligned sequences, facilitating comprehensive comparison and collinearity analysis with the T2T reference genome.

### Genome assembly with AutoHiC

During testing, all genomes only require the original genome files and Hi-C sequencing data. AutoHiC automatically performs error correction and generates result reports. The genome and interaction matrix files before and after correction are made available in the AutoHiC results for visualization and statistical analysis. The number of assembly errors can be directly obtained from the result report. Depending on the genome size, the Hi-C data size and the running time of AutoHiC for the number of correction iterations can be obtained from Supplementary Table 19.

### Implementation of the AtuoHiC framework

The deep learning architectures are implemented using MMDetection^53^ v.2.11.0 and PyTorch v.1.10.1 on Python v.3.7. Training is conducted on NVIDIA GeForce RTX 3080 Ti GPUs.

## Data availability

No new data were generated for this study. All data used in this article are available from Supplementary Table 17 and 18.

## Code availability

The AutoHiC source code and documentation are available on GitHub under the MIT license at https://github.com/Jwindler/AutoHiC.

## Acknowledgements

We thank Guoqing Zhang at Southwest University for critical reading of the manuscript. National Natural Science Foundation of China [U21A20248]; National Natural Science Foundation of China [32000340]; Fundamental Research Funds for the Central Universities [XDJK2019TJ003]. Funding for open access charge: National Natural Science Foundation of China.

## Author Contributions

Z.J. and Y.W. conceptualized the project. Y.W. supervised the project. Z.J. designed and wrote the software code. Y.L. completes the code portion of the results report. Z.J. and Z.P. conducted software testing and analysis. Z.J., Z.P. and L.B. performed data collection and labeling. Y.W. finalized the manuscript with input from all authors. All authors read and approved the final manuscript.

