## Supplementary Information for "AutoHiC: a deep-learning method for automatic and accurate chromosome-level genome assembly"

### Supplementary Notes

#### Supplementary Note 1. Scaffolder tool search.

To find all available (recent) Hi-C scaffolders, we performed a literature search in PubMed for publications between March 27, 2020 and March 27, 2023. The search terms "(hi-c scaffolding) or (hi-c assembly) or (hi-c genome assembly) or (hic scaffolding) or (hic assembly) or (hic genome assembly)" yielded 699 results. Nine scaffolding methods were identified, as well as the frequency with which they were used to publish genomes (Supplementary Table 1). We only keep 4 software, of which Lachesis no longer provides support, and the official website recommends using 3D-DNA instead. instaGRAAL can only run in the GPU environment. A software error occurred while running EndHiC. The author did not respond to the request. HiRise does not need to use Hi-C data. AllHiC is mainly used for plant genome assembly and requires genomes from homologous species.

#### Supplementary Note 2. Description of anomalous software assembly results.

In the comparison of the software, we found that the N50 values of Pin_hic and Yash are very large (Supplementary Table 3), basically close to the genome size, so we carefully studied their assembly results. We generated a histogram of the genome size and length of the top n sequences (n is the number of chromosomes) in the software assembly results (Supplementary Figures 9, 10, 11, 12, 13). It is obviously wrong to say that YaSH and Pin_hic merge multiple chromosomes together. To avoid misunderstanding, their results are shown in the appendix.

### Supplementary Figures

#### Summary

| | |
|---|---|
| Species | Brassica rapa |
| Assembly size (bp) | 262330601 |
| Scaffold N50 (bp) | 24553947 |
| Scaffold N90 (bp) | 19027318 |
| CC ratio (%) | 1.00 |
| Structural errors ratio (%) | 0 |
| Number of chromosomes | 11 |
| HiC Anchor rate (%) | 92.37 |
| Number of scaffolds | 11 |
| Longest scaffold (bp) | 34719908 |
| Scaffold L50 | 5 |
| Scaffold L90 | 10 |
| GC (%) | 35.09 |

**Supplementary Figure 1. AutoHiC Results Report Part I.**

The basic information of the genome assembled by AutoHiC, including genome size, N50, CC ratio, Anchor rate and GC content, etc.

### Adjusting result

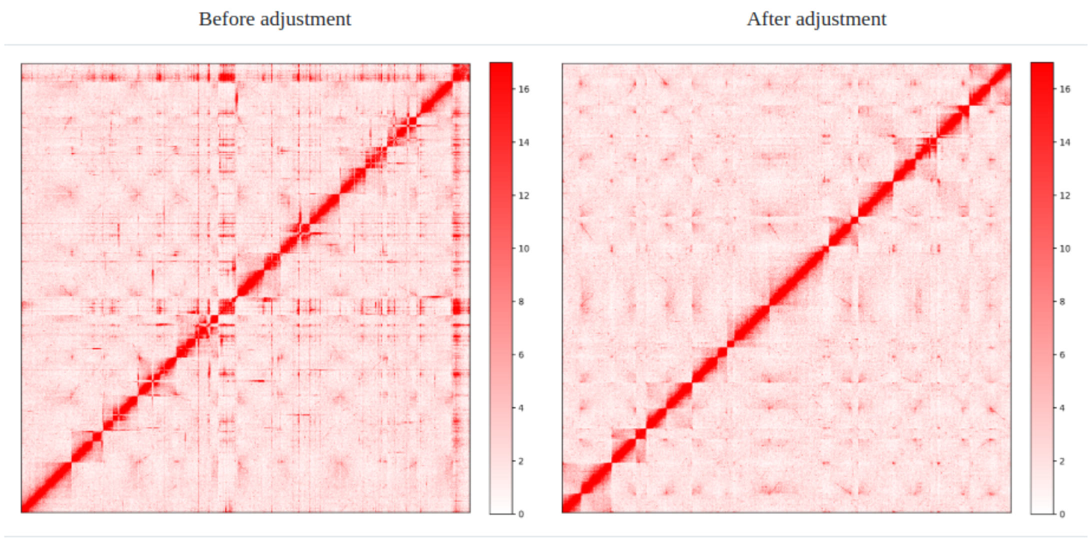


**Supplementary Figure 2. AutoHiC Results Report Part II.**

Comparison of global interaction heatmaps before and after AutoHiC error correction. Left is before adjustment. On the right is adjusted.

### Error adjustment

Translocation

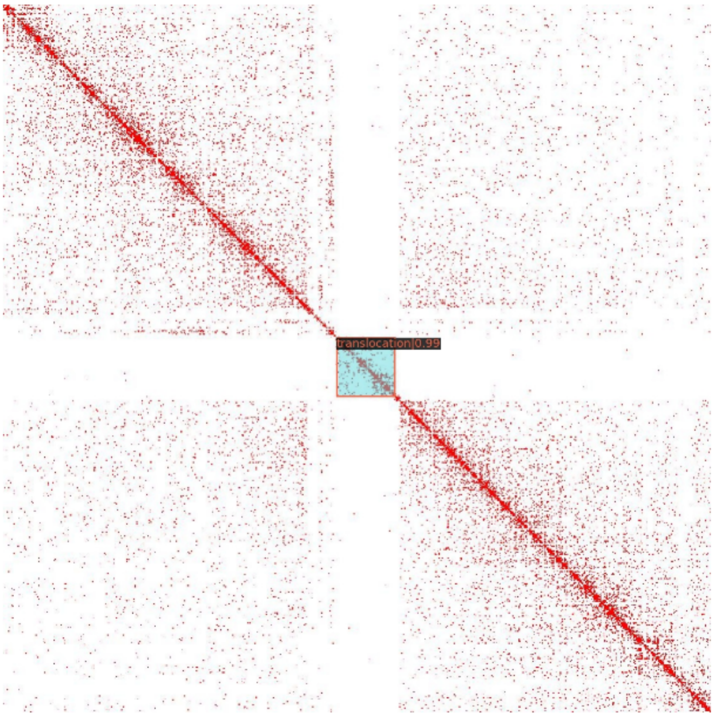


Location: 21648009-21934703

1

**Supplementary Figure 3. AutoHiC Results Report Part III.**

Error details detected by AutoHiC, which contains error visualization, location information.

### Chromosome length

| Molecule | Length(bp) | GC(%) |
|---|---|---|
| All | 262330601 | 35.09 |
| Chromosome 1 | 29096201 | 33.34 |
| Chromosome 2 | 20030698 | 33.37 |
| Chromosome 3 | 26915038 | 33.83 |
| Chromosome 4 | 20539817 | 33.91 |
| Chromosome 5 | 24553947 | 33.82 |
| Chromosome 6 | 34719908 | 34.28 |
| Chromosome 7 | 17287781 | 34.57 |
| Chromosome 8 | 20321949 | 33.79 |
| Chromosome 9 | 25536935 | 33.99 |
| Chromosome 10 | 19027318 | 33.13 |
| Chromosome 11 | 24301009 | 33.79 |

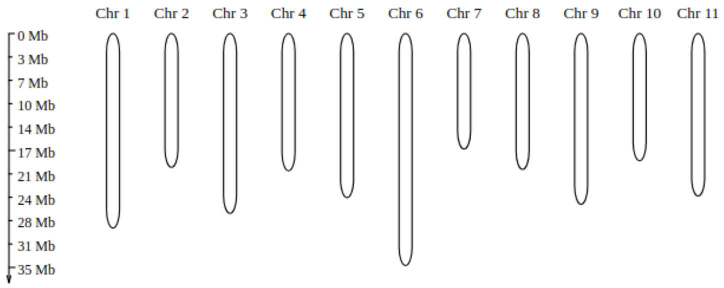


**Supplementary Figure 4. AutoHiC Results Report Part IV.**

Chromosome number and length predicted by AutoHiC, and calculated GC content of each chromosome. Below is a visualization of the chromosomes.

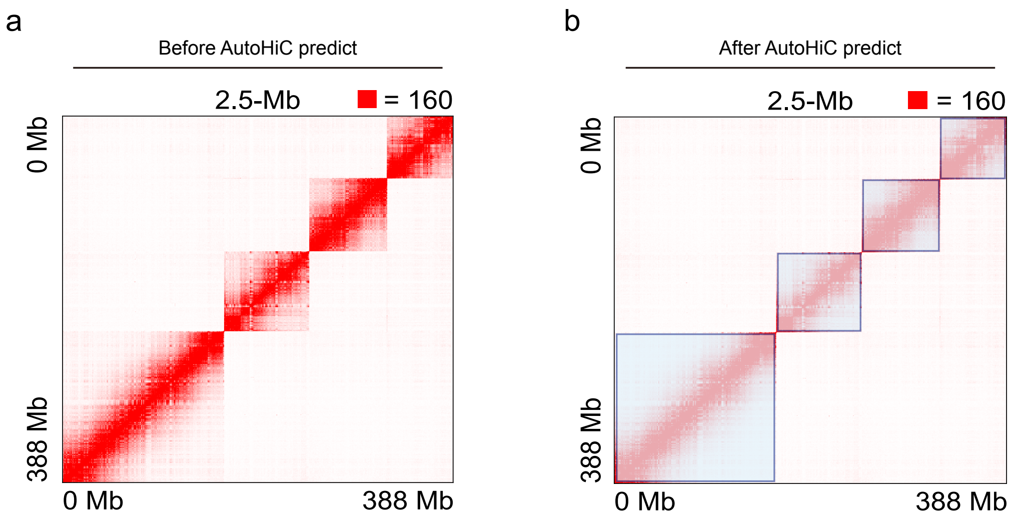


**Supplementary Figure 5. AutoHiC predicts chromosome number and length.**

**a** Chromosome global interaction heatmap before AutoHiC prediction. **b** Chromosome global interaction heatmap after AutoHiC prediction. The area surrounded by the blue box is the interaction area within the chromosome..

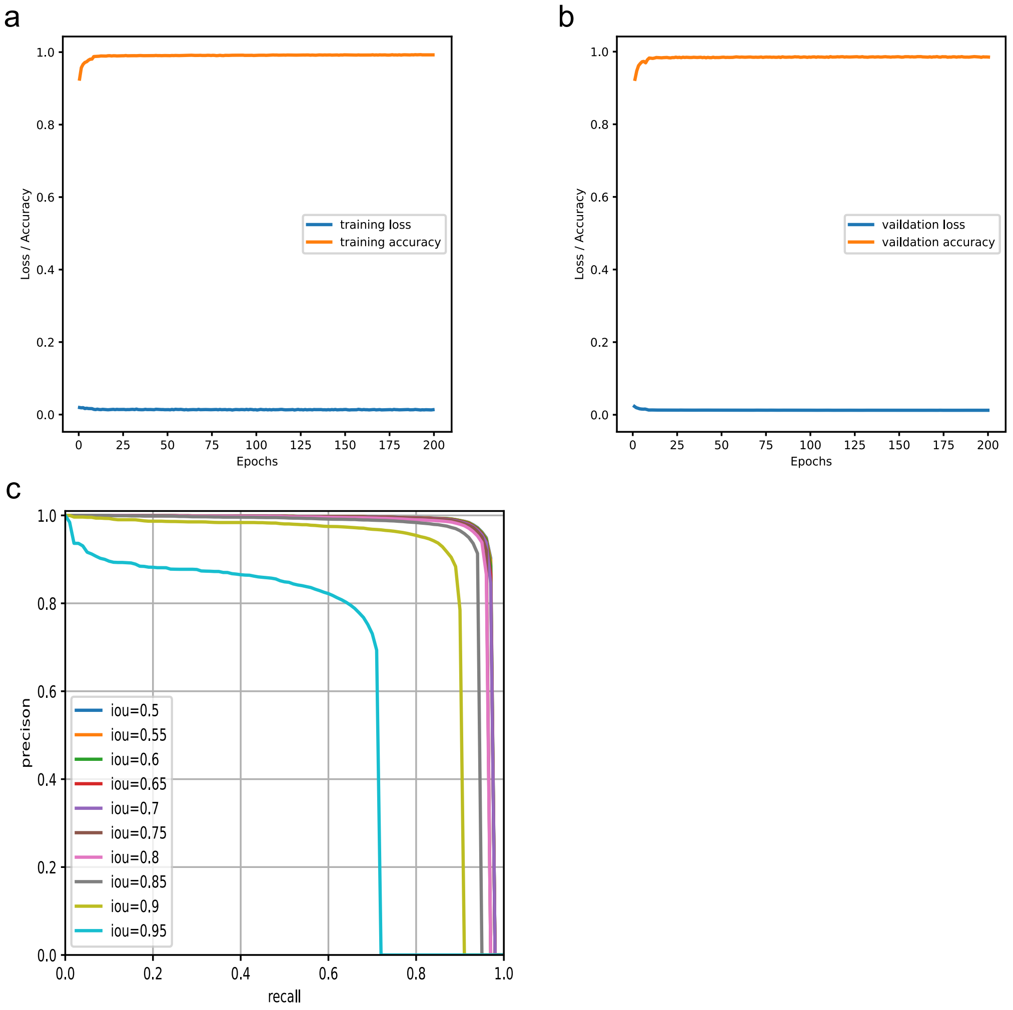


**Supplementary Figure 6. Chromosome identification model evaluation.**

**a**, **b** Changes in model accuracy and loss rate during training and validation. **c** Precision Recall (PR) curve for the different IOU thresholds as in panel.

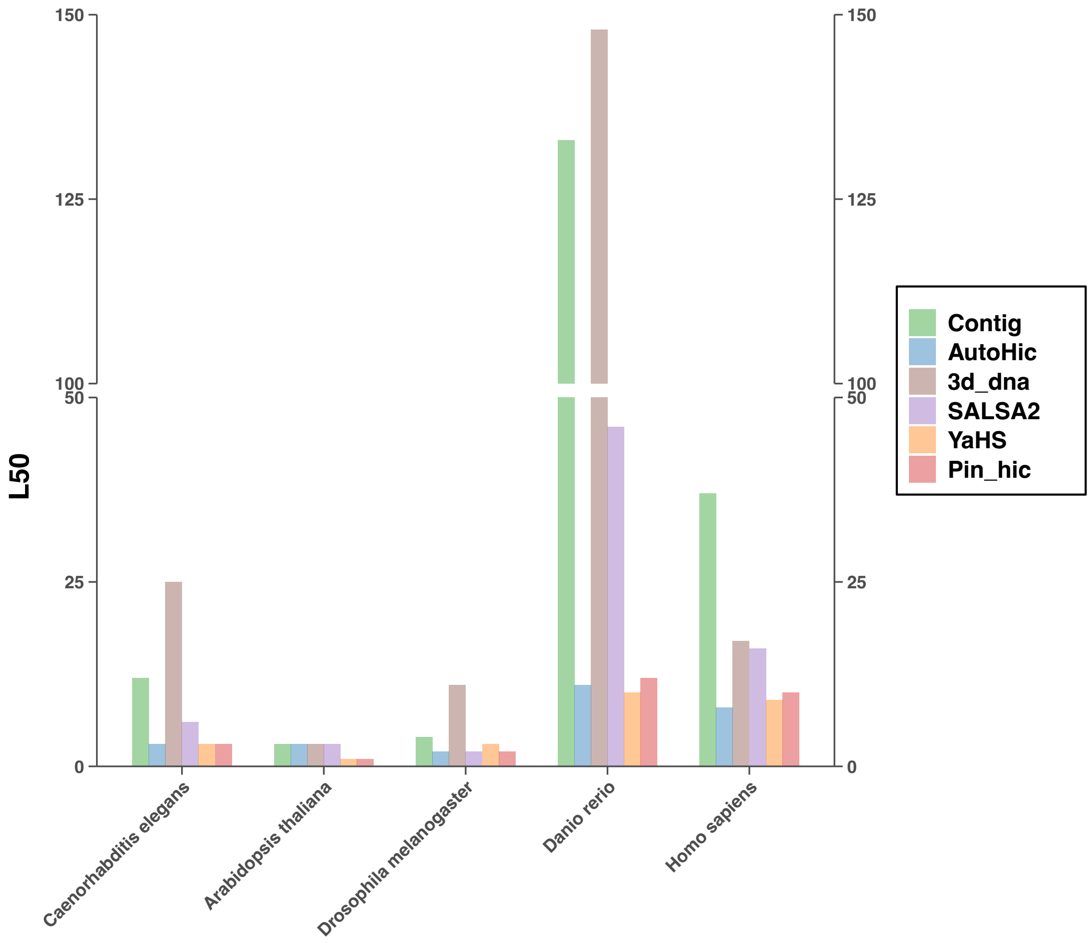


**Supplementary Figure 7. Performance evaluation of L50 on the five species.**

Histogram of L50 values for different software contig results. It contains contig results, and different colors represent different software.

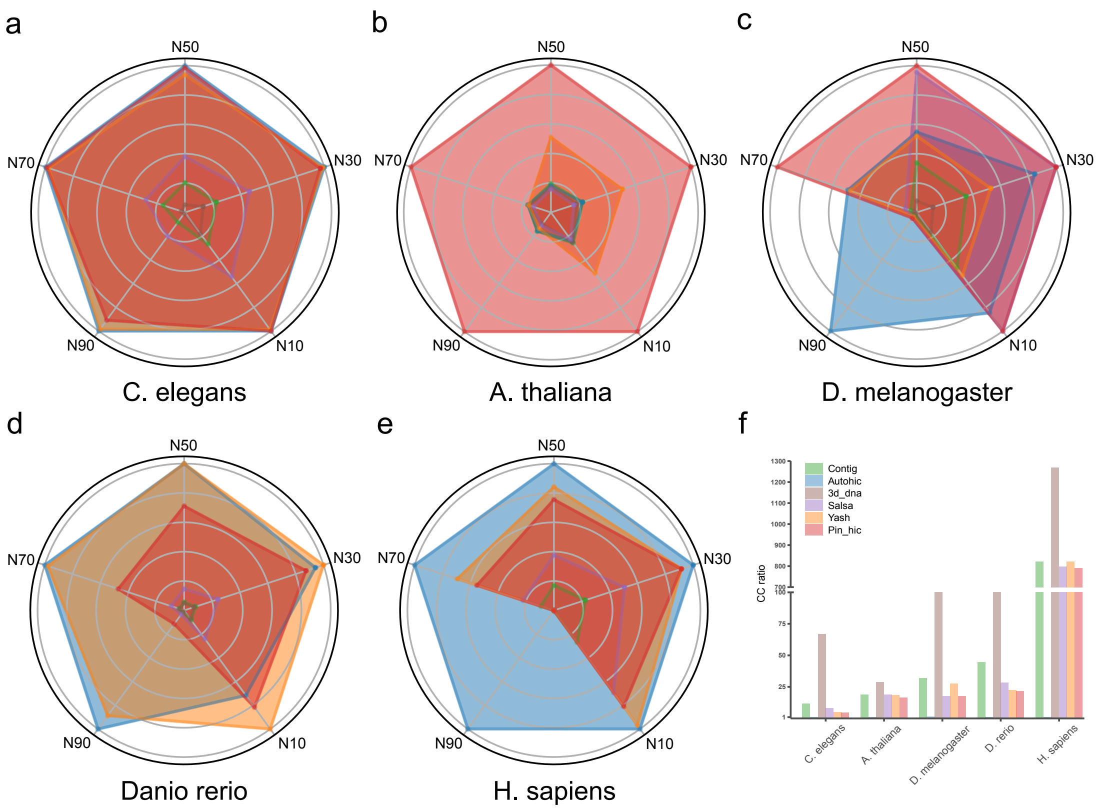


**Supplementary Figure 8. Performance evaluation of continuity on the five species.**

**a**, **b**, **c**, **d**, **e** The radar chart shows the continuity of the assembly results. Different colors represent different software. **f** Histogram of CC rate values of different software contig results. It contains the results of the contig, and different colors represent different software.

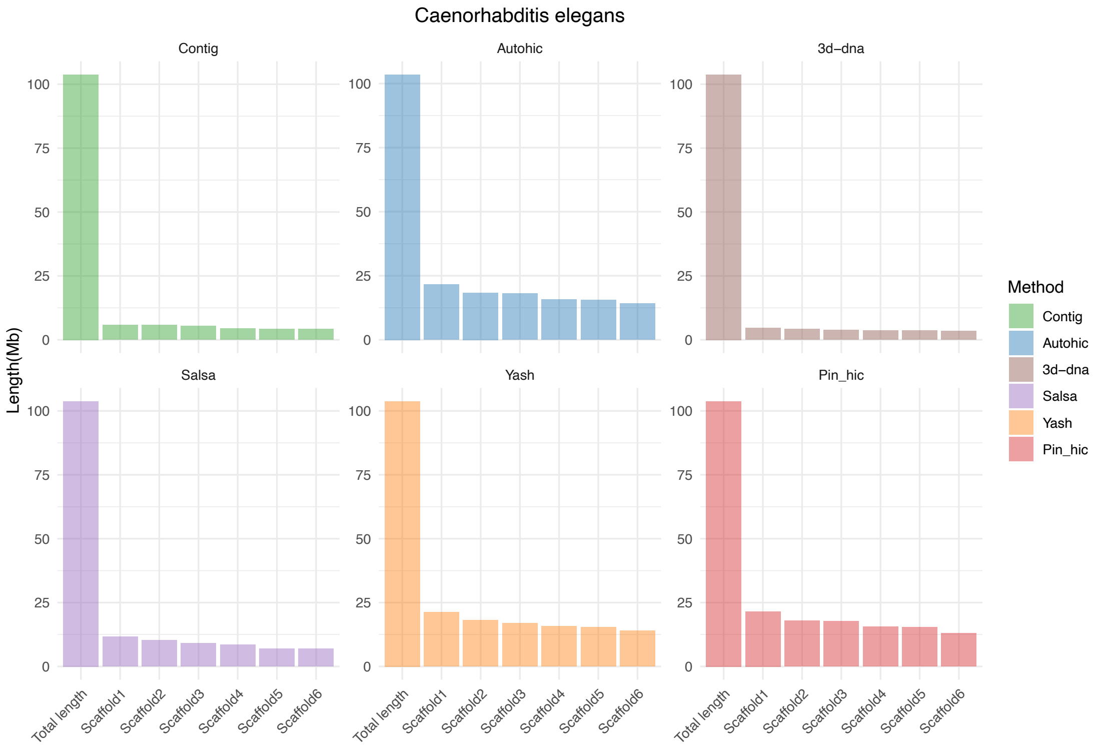


**Supplementary Figure 9. Genome and scaffold length of Caenorhabditis elegans.**

Only the genome and the length of the first n scaffolds are counted, where n is the number of chromosomes. One panel per software.

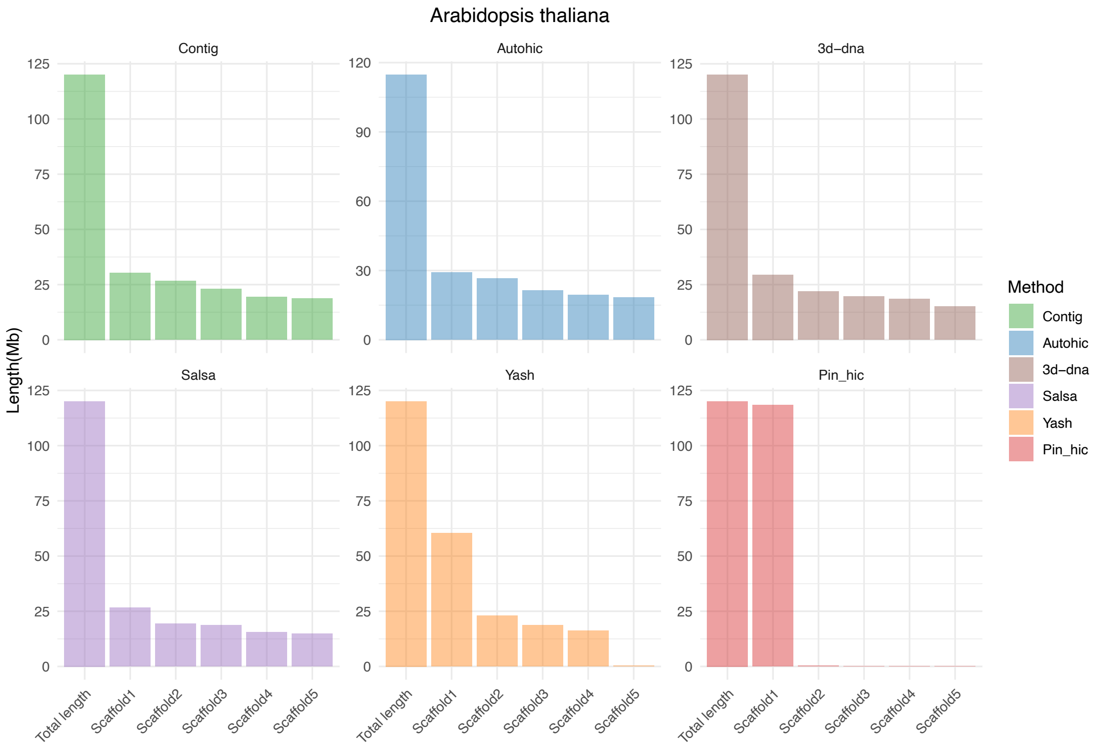


**Supplementary Figure 10. Genome and scaffold length of *Arabidopsis thaliana*.**

Only the genome and the length of the first n scaffolds are counted, where n is the number of chromosomes. One panel per software.

.

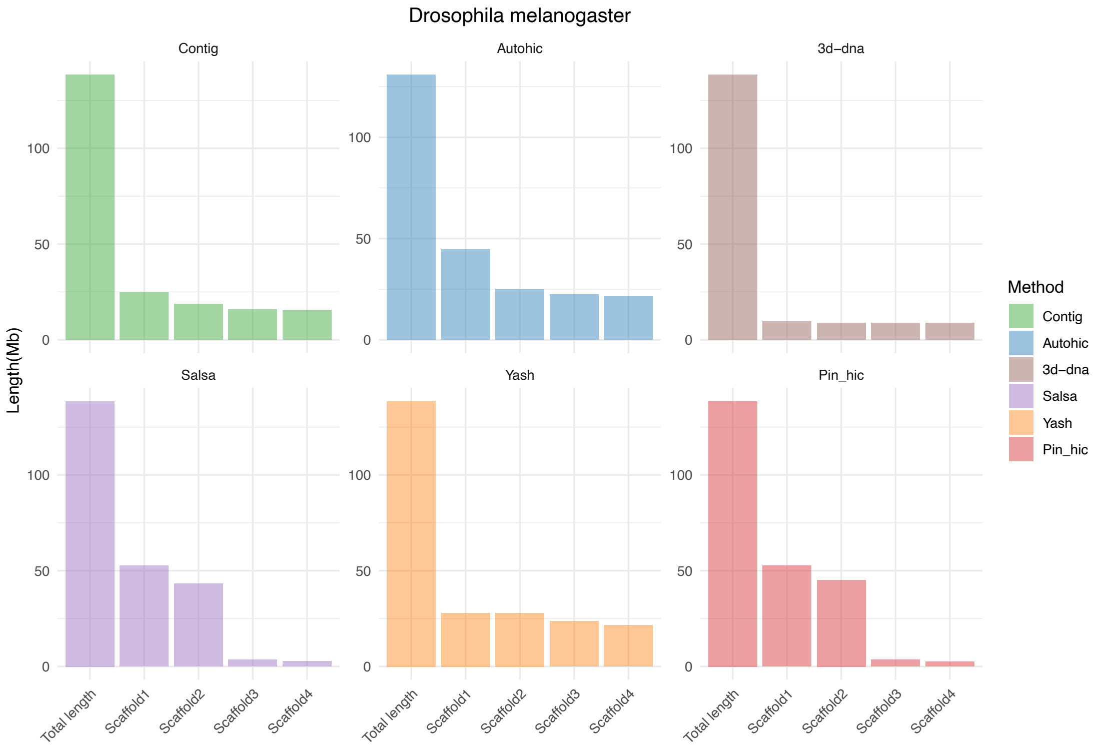


**Supplementary Figure 11. Genome and scaffold length of *Drosophila melanogaster*.** Only the genome and the length of the first n scaffolds are counted, where n is the number of chromosomes. One panel per software.

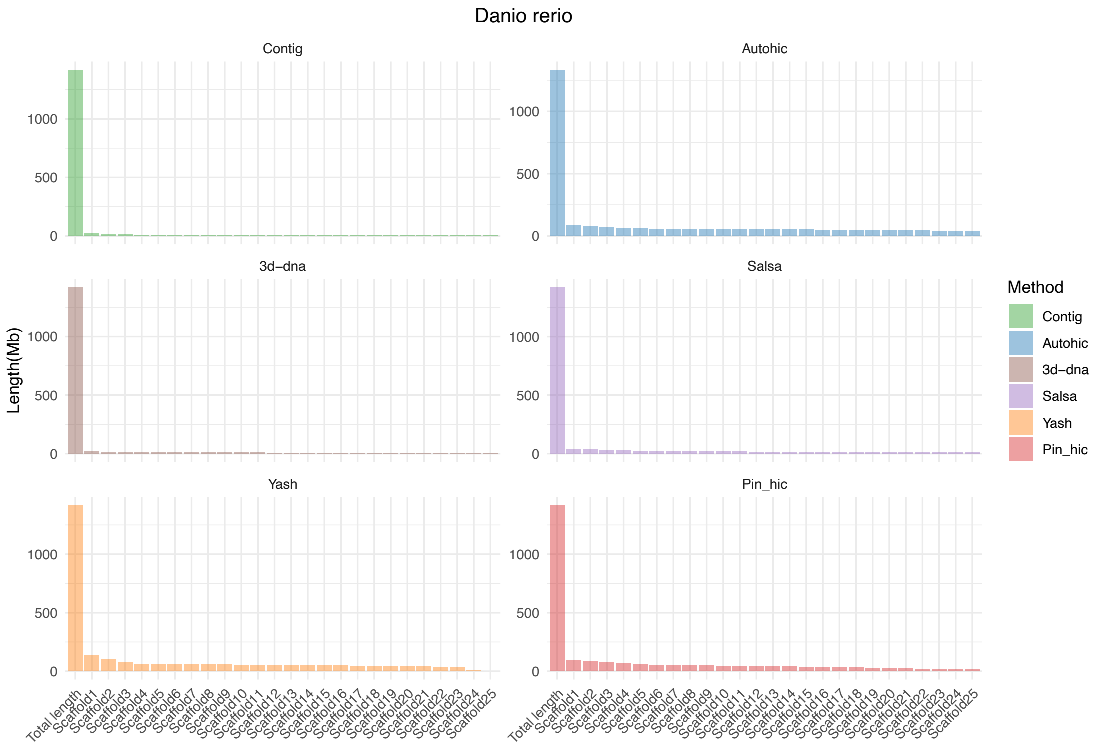


**Supplementary Figure 12. Genome and scaffold length of *Danio rerio*.**

Only the genome and the length of the first n scaffolds are counted, where n is the number of chromosomes. One panel per software.

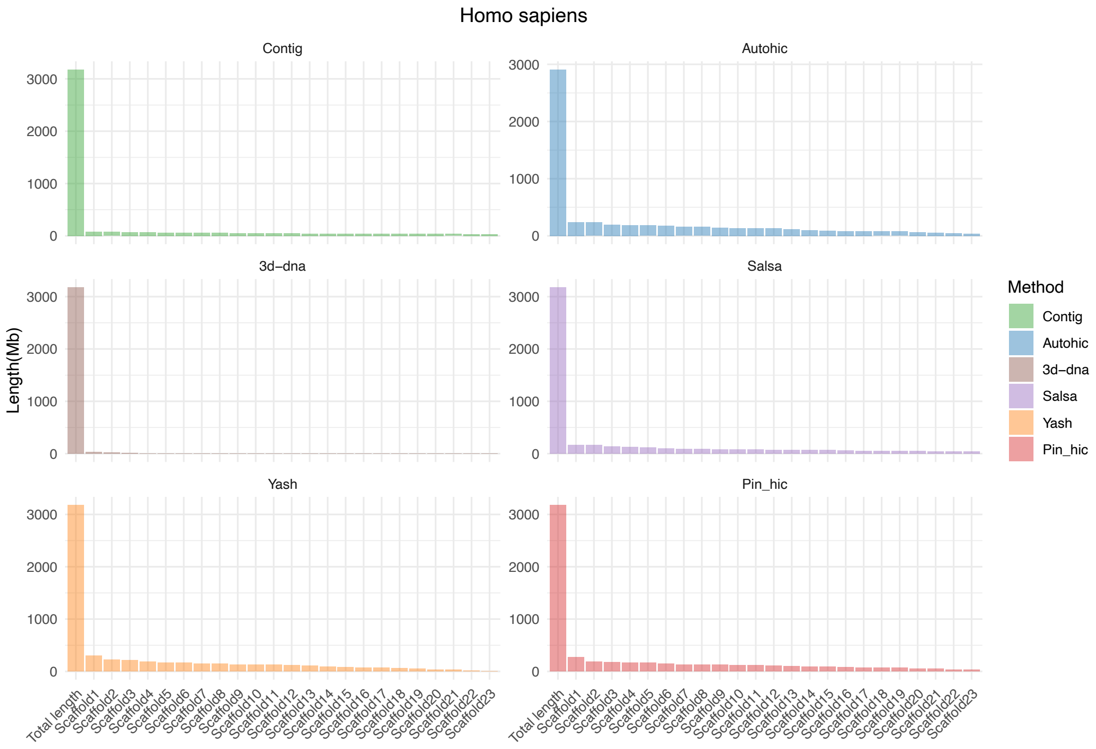


**Supplementary Figure 13. Genome and scaffold length of *Homo sapiens*.**

Only the genome and the length of the first n scaffolds are counted, where n is the number of chromosomes. One panel per software.

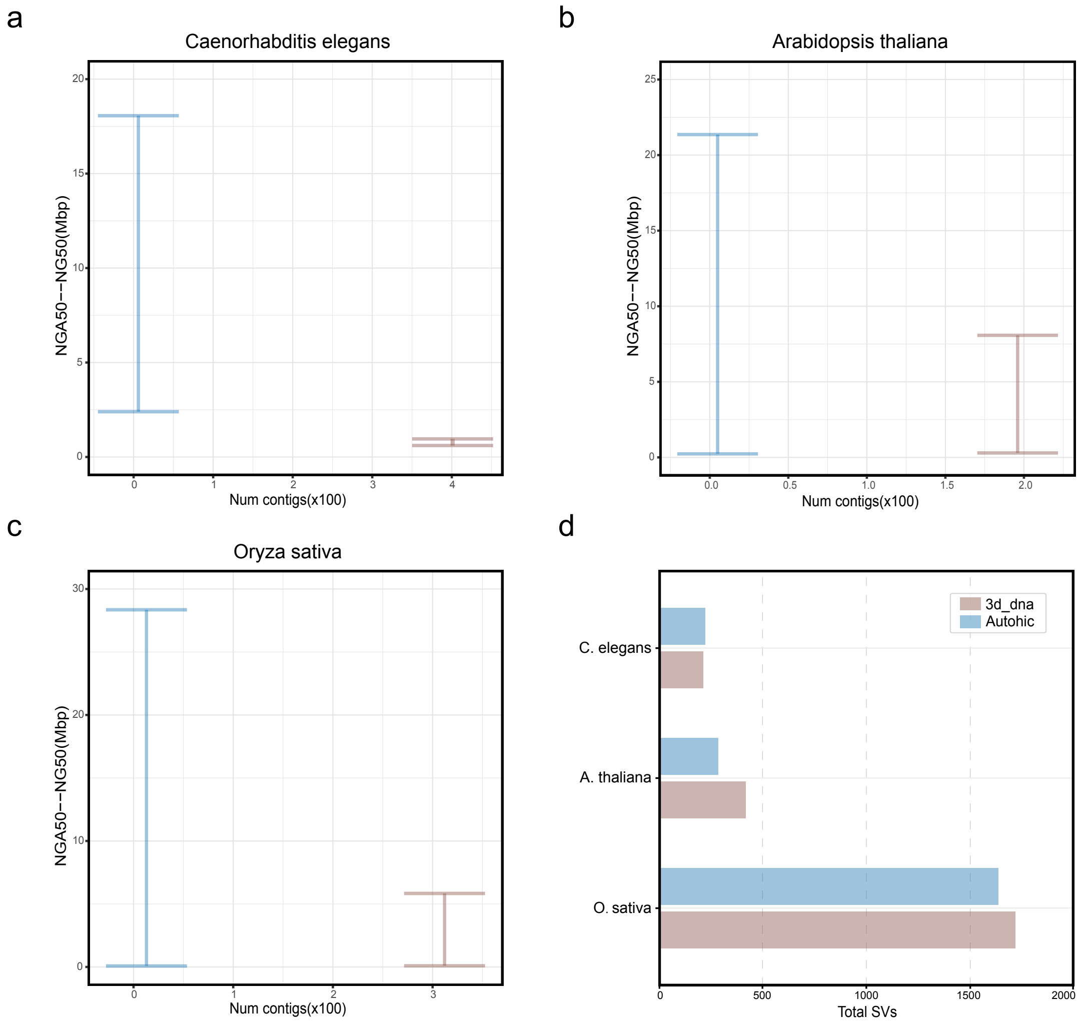


**Supplementary Figure 14. Assessing the performance of correcting the T2T genome.**
**a, b, c** Comparison of the contiguity of assemblies scaffolded by 3D-DNA and AutoHiC as measured by QUAST. The Y-axes show the range of NGA50 to NG50 lengths to indicate the uncertainty caused by true genomic variation between the individual and the reference genome. **d** Comparison of the accuracy of assemblies scaffolded by 3D-DNA and AutoHiC as measured by MUM&Co. The X-axis shows the total number of structural variations to show AutoHiC's ability to correct errors.

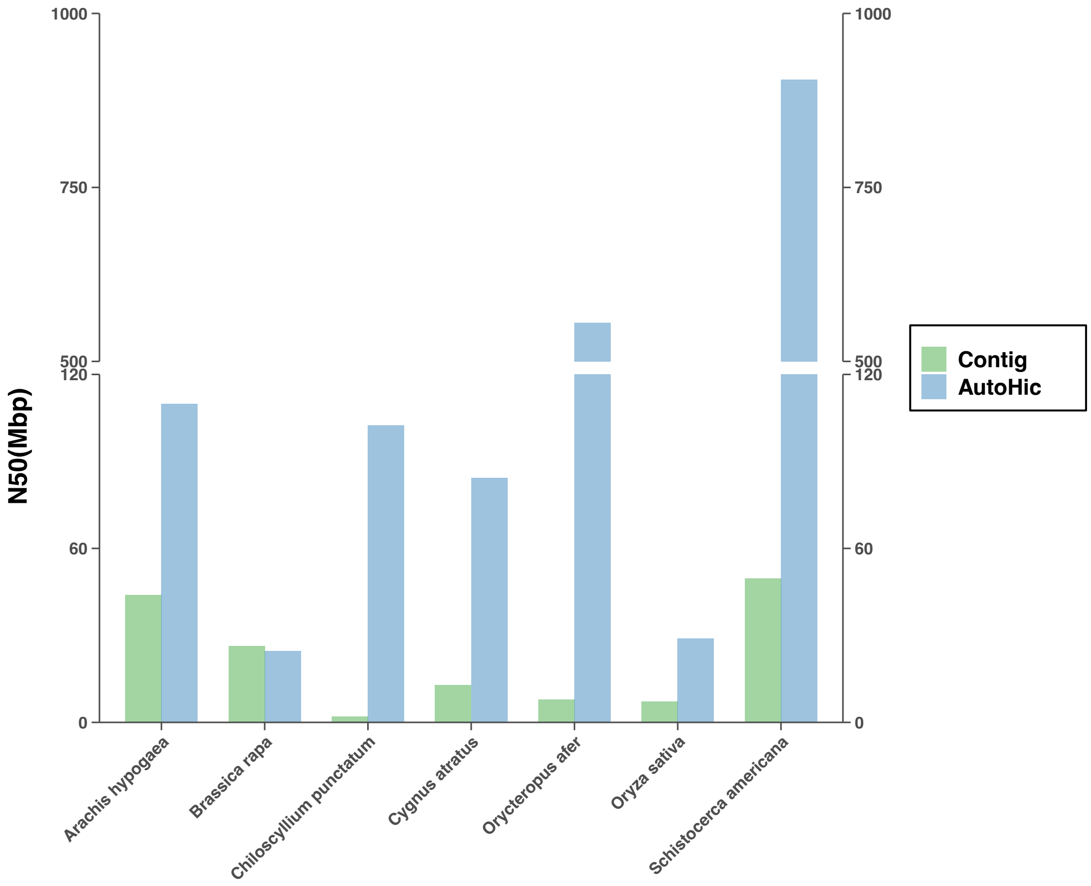


**Supplementary Figure 15. N50 histogram before and after genome assembly.**

Changes in N50 before and after genome assembly with AutoHiC. The Y-axis is the size of N50 and the X-axis is the different genomes. The colors represent the genome before and after assembly.

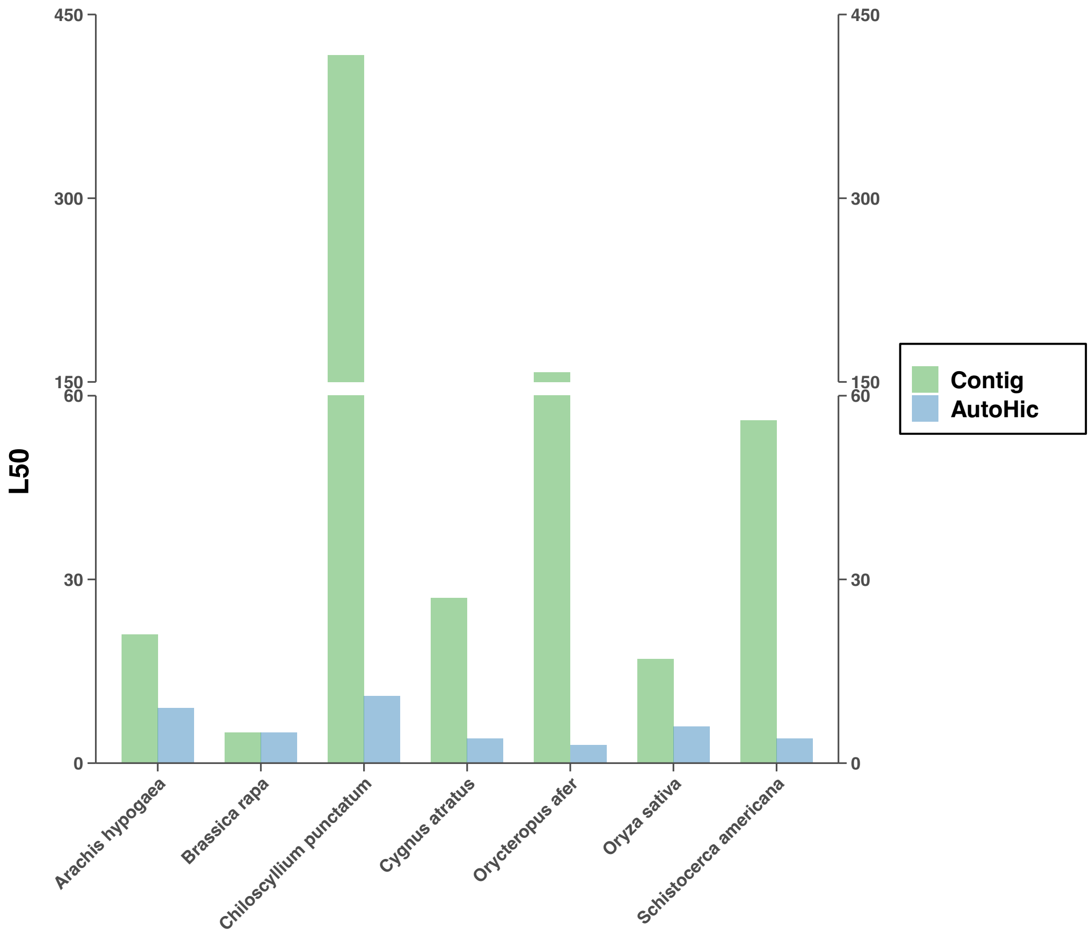


**Supplementary Figure 16. L50 histogram before and after genome assembly.**

Changes in L50 before and after genome assembly with AutoHiC. The Y-axis is the size of L50 and the X-axis is the different genomes. The colors represent the genome before and after assembly.

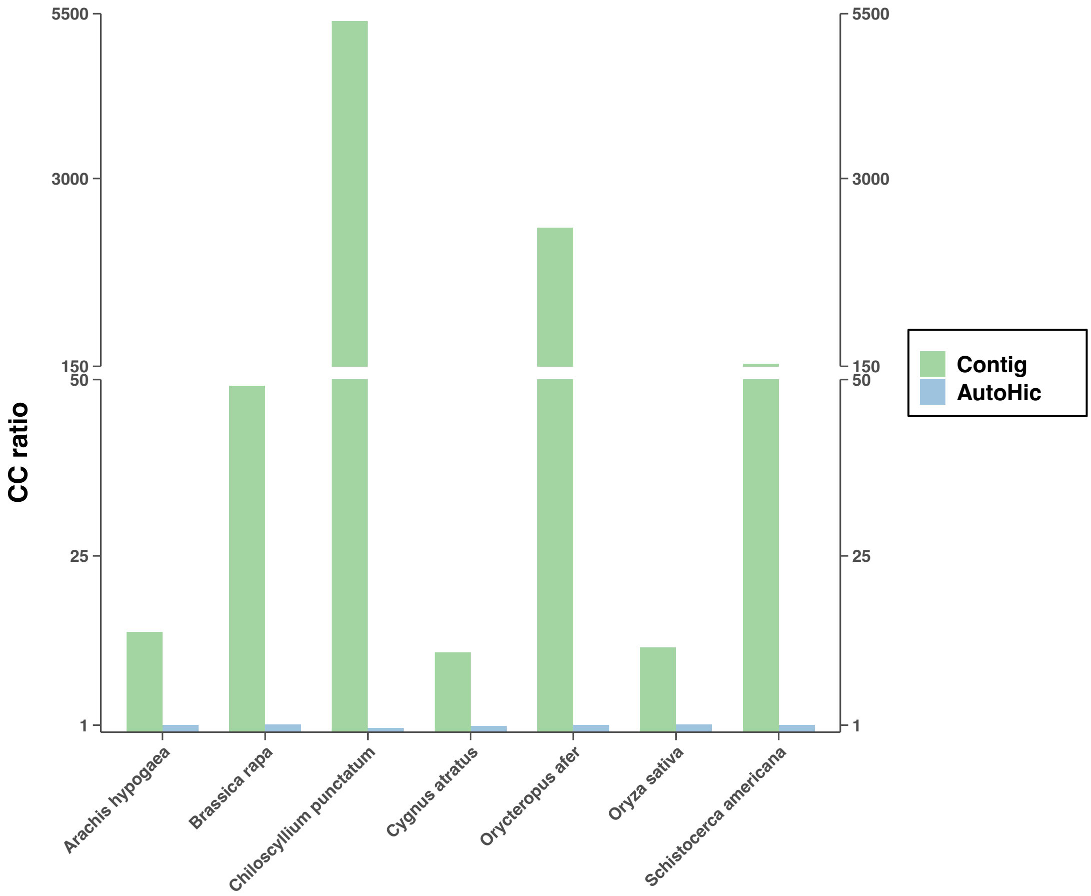


**Supplementary Figure 17. CC ratio histogram before and after genome assembly.** Changes in CC ratio before and after genome assembly with AutoHiC. The Y-axis is the size of CC ratio and the X-axis is the different genomes. The colors represent the genome before and after assembly.

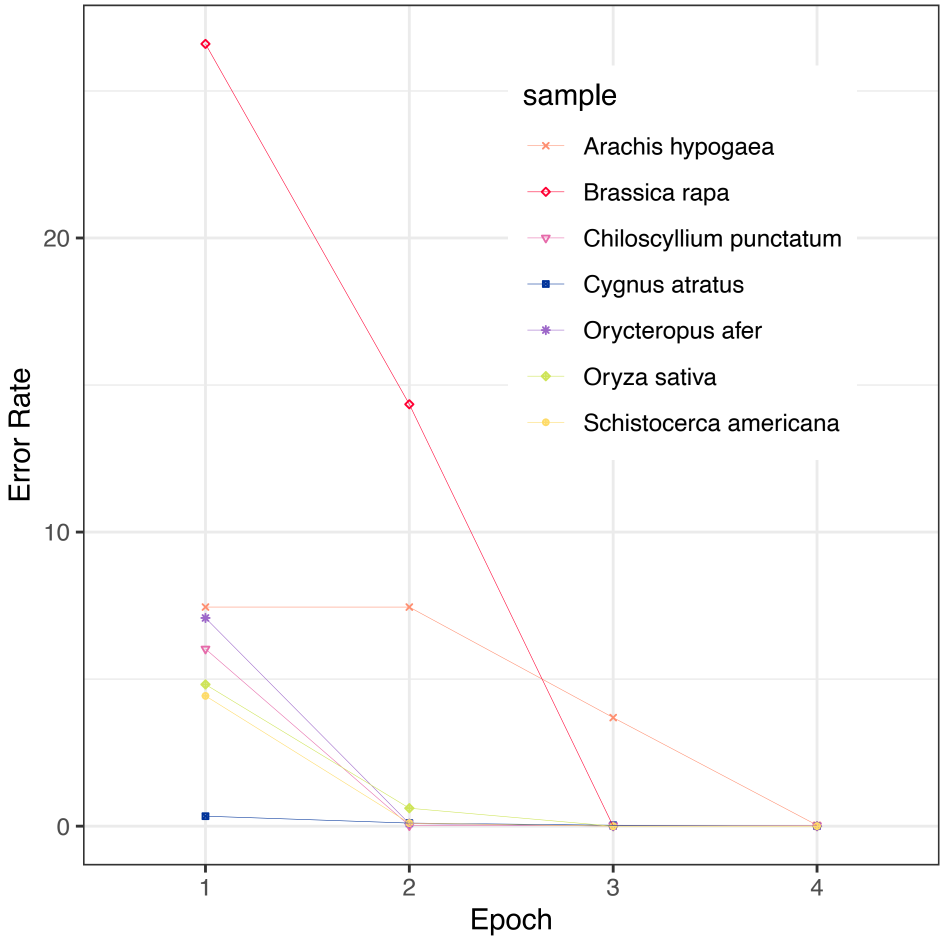


**Supplementary Figure 18. Error rate variation curve during genome assembly.**

Variation of the error rate for assembly errors corrected by AutoHiC. The X-axis is the number of iterative error corrections. The Y-axis is the magnitude of the error rate, scaled 1000 folds.

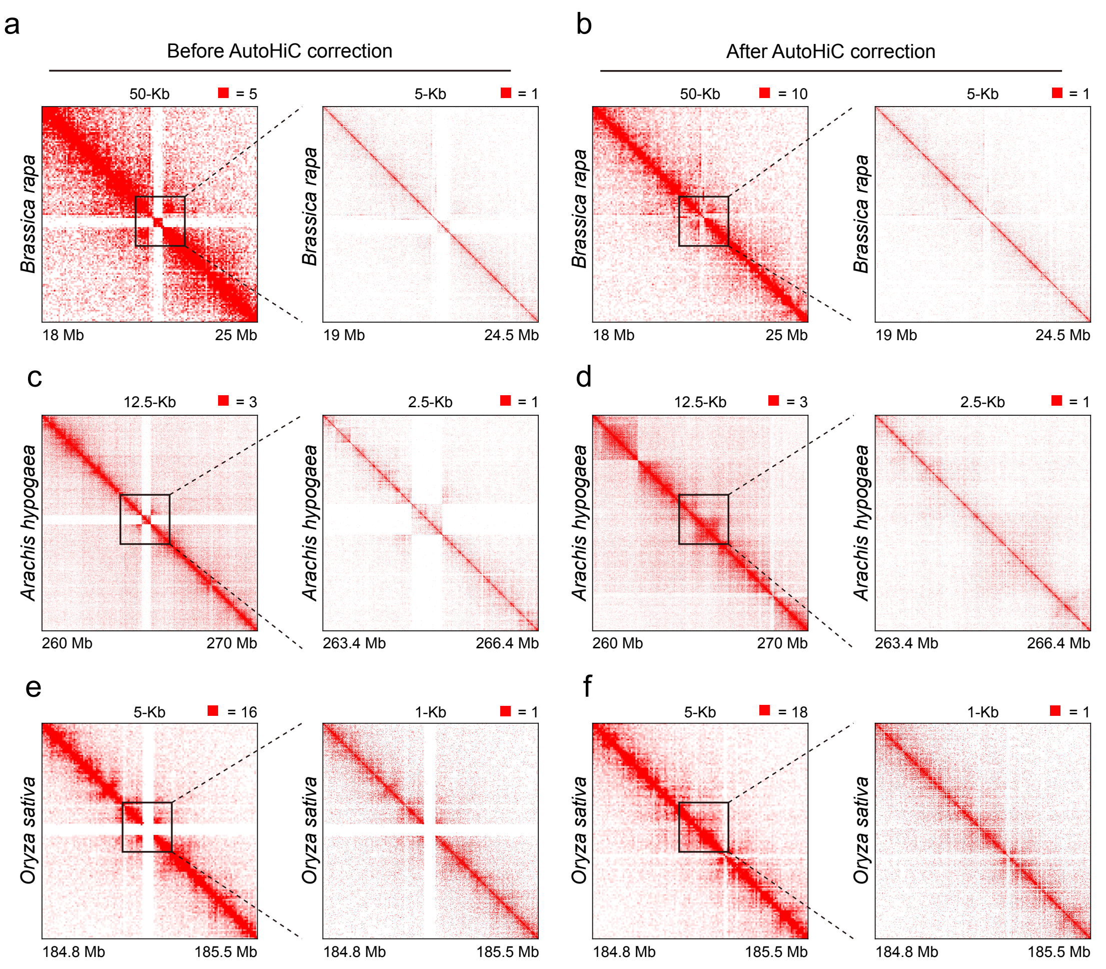


**Supplementary Figure 19. Compare interaction heatmaps before and after AutoHiC correction.**

**a**, **b** Heatmap of interactions before and after translocation error regions during correcting in *Brassica rapa*. **c**, **d** Heatmap of interactions before and after translocation error regions during correcting in *Arachis hypogaea*. **e**, **f** Heatmap of interactions before and after debris error regions during correcting in *Oryza sativa*. Low resolution on the left, high resolution on the right.

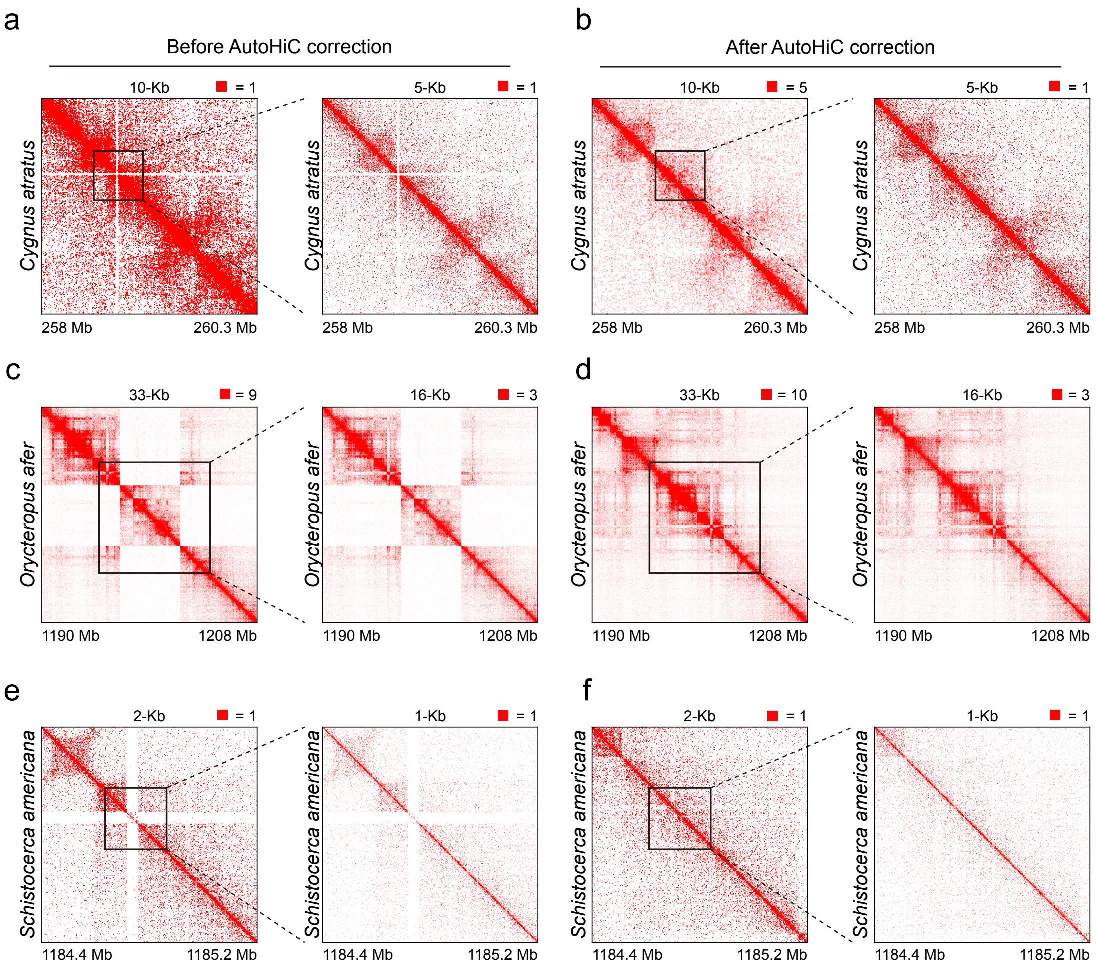


**Supplementary Figure 20. Compare interaction heatmaps before and after AutoHiC correction.**

**a**, **b** Heatmap of interactions before and after translocation error regions during correcting in *Cygnus atratus*. **c**, **d** Heatmap of interactions before and after translocation error regions during correcting in *Orycteropus afer*. **e**, **f** Heatmap of interactions before and after translocation error regions during correcting in *Schistocerca americana*. Low resolution on the left, high resolution on the right.

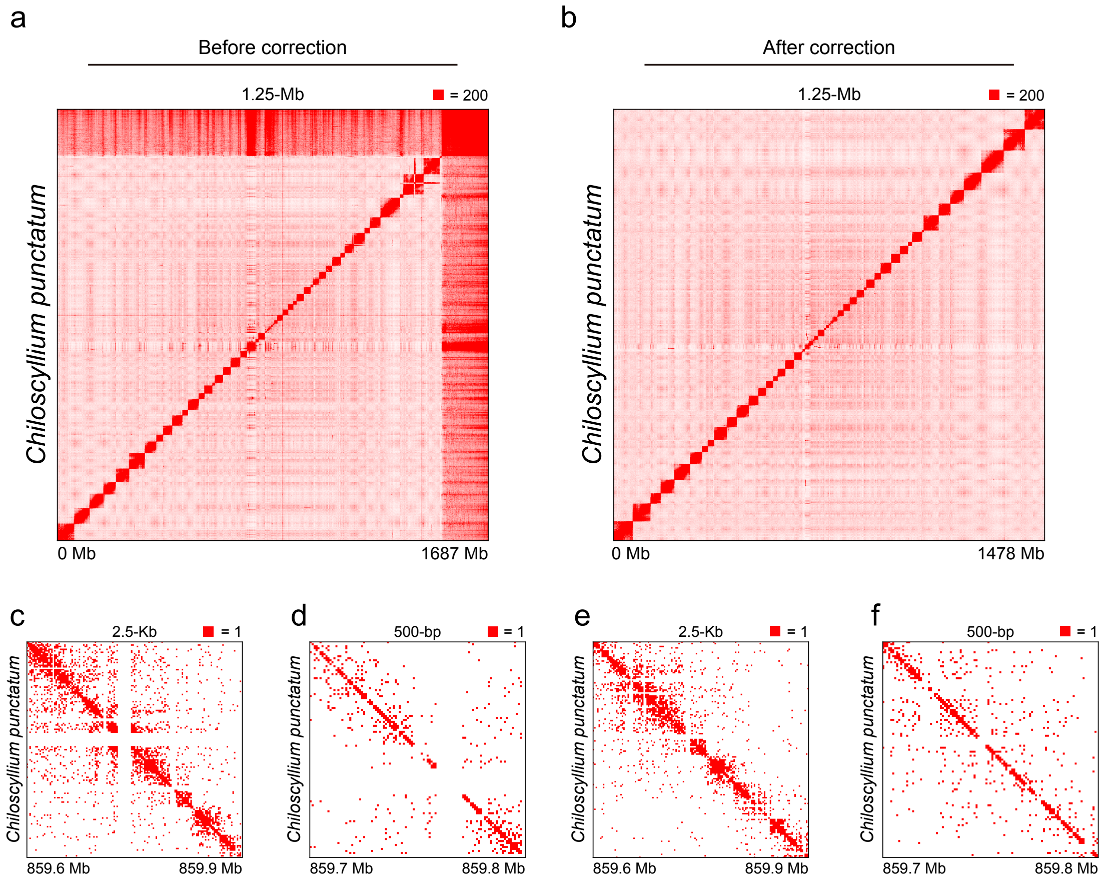


**Supplementary Figure 21. Compare interaction heatmaps before and after AutoHiC correction.**

**a**, **b** Comparison of global interaction heatmaps of *Chiloscyllium punctatum* before and after fitting. Left is before adjustment. Right is after fitting. **c**, **d, e**, **f** Heatmap of interactions before and after debris error regions during correcting in *Chiloscyllium punctatum*. Low resolution on the left, high resolution on the right.

### Supplementary Tables

**Supplementary Table 1. scaffolding software statistics.**

| Name | Publication Date | Citation Count | Application Count |
|---|---|---|---|
| Lachesis | December 1, 2013 | 860 | 271 |
| 3D-DNA | Mar 23, 2017 | 808 | 256 |
| SALSA2 | August 21, 2019 | 363 | 68 |
| yahs | December 16, 2022 | 78 | 5 |
| instaGRAAL | June 1, 2020 | 24 | 4 |
| EndHiC | December 8, 2022 | 3 | 1 |
| Pin_hic | December 31, 2021 | 0 | 0 |
| HiRise | March 26, 2016 | 604 | 180 |
| AllHiC | August 5, 2019 | 176 | 47 |
| SALSA1 | July 12, 2017 | 104 | 54 |

**Supplementary Table 2. QUAST results comparing software assembly results.**

Please see separate table file.

**Supplementary Table 3. Nx index statistics for comparing software assembly results.**

Please see separate table file.

**Supplementary Table 4. Assembly result sequence length statistics.**

Please see separate table file.

**Supplementary Table 5. QUAST results of T2T assembly results.**

Please see separate table file.

**Supplementary Table 6. *Homo sapiens* MUM&Co results before correcting errors.**

Please see separate table file.

**Supplementary Table 7.** ***Homo sapiens*** **MUM&Co results after correcting errors.**

Please see separate table file.

**Supplementary Table 8.** ***Bombyx mori*** **MUM&Co results before correcting errors.**

Please see separate table file.

**Supplementary Table 9.** ***Bombyx mori*** **MUM&Co results after correcting errors.**

Please see separate table file.

**Supplementary Table 10.** ***Caenorhabditis elegans*** **MUM&Co results before correcting errors.**

Please see separate table file.

**Supplementary Table 11.** ***Caenorhabditis elegans*** **MUM&Co results after correcting errors.**

Please see separate table file.

**Supplementary Table 12.** ***Arabidopsis thaliana*** **MUM&Co results before correcting errors.**

Please see separate table file.

**Supplementary Table 13.** ***Arabidopsis thaliana*** **MUM&Co results after correcting errors.**

Please see separate table file.

**Supplementary Table 14.** ***Oryza sativa*** **MUM&Co results before correcting errors.**

Please see separate table file.

**Supplementary Table 15.** ***Oryza sativa*** **MUM&Co results after correcting errors.**

Please see separate table file.

**Supplementary Table 16. QUAST results on the extended test dataset.**

Please see separate table file.

**Supplementary Table 17. Genome data and Hi-C raw data statistics.**

Please see separate table file.

**Supplementary Table 18. Model Training Dataset Statistics.**

Please see separate table file.

**Supplementary Table 19. AutoHiC runtime.**

Please see separate table file.
